# p75 neurotrophin receptor signaling through the RhoA/ROCK pathway contributes to Alzheimer’s Disease tauopathy

**DOI:** 10.64898/2026.02.22.707328

**Authors:** Kang Liao, Meng Xie, Carlos F. Ibáñez

**Author notes:** Corresponding author: C.F.I.

## Abstract

Therapeutic development in Alzheimer’s Disease (AD) has for the most part been focused on reducing β-amyloid load. Nevertheless, neurofibrillary tangles (NFTs), produced by aggregation of hyper-phosphorylated Tau protein, correlate with neurodegeneration and cognitive impairment significantly better than amyloid accumulation in AD patients. Here we report that P301S mice, a model of AD tauopathy, carrying mutant variants of the p75 neurotrophin receptor (p75^NTR^) deficient in RhoA/ROCK signaling are protected from neurodegeneration and cognitive impairment. Both p75^ΔDD^, lacking the death domain, and triple mutant p75^KKEA^, unable to interact with RhoGDI, decreased NFT levels, reduced gliosis, neurodegeneration and synapse loss, and improved spatial learning and memory in P301S mice. Intriguingly, p75^C259A^, a variant unresponsive to neurotrophins but still competent for RhoA signaling induced by myelin-derived ligands, did not afford any neuroprotection. P301S neurons expressing p75^ΔDD^ or p75^KKEA^, but not p75^C259A^, showed reduced phospho-Tau and ROCK and GSK3β activity, the two main kinases responsible for Tau phosphorylation. In line with this, treatment with myelin-associated glycoprotein (MAG) enhanced Tau phosphorylation and ROCK activity in P301S neurons expressing wild type p75^NTR^ or p75^C259A^, but not p75^ΔDD^ or p75^KKEA^. Together, these results indicate that p75^NTR^ contributes to AD tauopathy by enhancing the activity of the RhoA-ROCK pathway.

## Introduction

Two key features characterize the histopathology of Alzheimer’s Disease (AD), namely extracellular deposition of amyloid plaques, produced by aggregation of the amyloid β (Aβ) peptide, and intracellular neurofibrillary tangles (NFTs), fibrillar aggregates of hyper-phosphorylated Tau protein (Long and Holtzman, 2019). Familial cases of AD (FAD), approximately 5% of all AD cases, are characterized by mutations in the genes encoding Amyloid Precursor Protein (APP), from which the Aβ peptide is generated, and presenilin (PSEN), a critical component of γ-secretase, the key protease that generates Aβ (Beyreuther and Masters, 1991; Selkoe, 1991; Hardy and Allsop, 1991). The study of FAD cases has established a principal causative link between Aβ production and AD pathogenesis, also known as the “amyloid hypothesis” (Selkoe and Hardy, 2016). Perhaps because of this, most efforts regarding therapeutic development in AD have been devoted to counteract production or enhance clearance of Aβ and the build up of amyloid (Zhang et al., 2024; Manap et al., 2024). Recently, and for the first time, two anti-β amyloid antibodies, i.e. lecanemab and donanemab, have been approved for the treatment of AD in US, EU, China and several other countries after showing promising results in clinical trials (Fox et al., 2025). In contrast, Tau and NFTs have attracted much less attention in clinical development. A recent review listed 32 completed or ongoing clinical trials based on β amyloid and only 5 on Tau-related strategies (Manap et al., 2024). This is despite the fact that NFT histopathology and hyper-phosphorylated Tau correlate with neurodegeneration and cognitive impairment significantly better than Aβ accumulation in AD patients [Braak 1991; Arboleda-Velasquez 2019; Josephs 2008; Lopera 2023]. Other current clinical approaches in AD include strategies addressing neuroinflammation, neuroprotection and metabolism (Frisoni et al., 2025; Zhang et al., 2024; Manap et al., 2024). Clearly, a therapeutic strategy that simultaneously targets amyloid β and Tau NFTs would be most desirable.

Unlike FAD patients, mouse models of AD based on FAD mutations in APP or PSEN only present β amyloid pathology; neither hyper-phosphorylated Tau nor NFTs are present in these mice (Wilcock et al., 2008; Pang et al., 2021). For this reason, AD tauopathy has been modeled in mice through expression of human Tau carrying mutations associated with frontotemporal dementia such as P301L and P301S (Allen et al., 2002; Deters et al., 2008). These mice present Tau hyper-phosphorylation and NFTs, followed by neurodegeneration, neuroinflammation and learning and memory impairments, thus recapitulating key features of human tauopathy (Yoshiyama et al., 2007). Among the kinases that hyper-phosphorylate Tau in these models as well as AD patients are Rho-associated kinase (ROCK) and Glycogen Synthase Kinase-3 beta (GSK3β), the latter also a downstream target of the RhoA/ROCK pathway (Medd et al., 2025; Amano et al., 2003). In line with this, inhibitors of ROCK have been shown to reduce Tau hyper-phosphorylation and aggregation (Hamano et al., 2020; Zheng et al., 2025). Unfortunately, given the multiple, pleiotropic effects of ROCK in cell physiology, the current inhibitors are expected to have numerous unwanted effects and can therefore not be used in the clinic.

The p75 neurotrophin receptor (p75^NTR^), a member of the death receptor superfamily characterized by a death domain in its intracellular region, initiates diverse signaling cascades in response to neurotrophins, such as nerve growth factor (NGF), as well as other ligands, including myelin components such as myelin-associated glycoprotein (MAG) (Ibáñez and Simi, 2012; Underwood and Coulson, 2008). Three major signaling pathways can be regulated by p75^NTR^ in response to ligand binding, namely NF-kB signaling, c-Jun kinase (JNK) signaling followed by caspase-3 activation, and RhoA signaling leading to ROCK activation (Ibáñez and Simi, 2012; Underwood and Coulson, 2008). p75^NTR^ expression becomes upregulated following neural injury and cellular stress, including AD (Chakravarthy et al., 2012; Hu et al., 2002; Mufson and Kordower, 1992; Ernfors et al., 1990), and its activation can contribute to neural death, axonal degeneration and synaptic dysfunction (reviewed in (Ibáñez and Simi, 2012)). In line with this, ablation of p75^NTR^ signaling or expression reduces AD pathology and improves cognitive functions in mouse models of AD based on APP mutations (Knowles et al., 2009; Wang et al., 2011; Yi et al., 2021), as well as the Tau P301L mutation (Mañucat-Tan et al., 2019; Shen et al., 2018). In our earlier work, we took advantage of signaling-deficient p75^NTR^ mutants lacking either the death domain (ΔDD) or transmembrane Cys^259^ (C259A), both of which are essential for receptor activity in response to neurotrophins, to uncover a previously unknown role for p75^NTR^ in APP internalization, amyloidogenic processing and Aβ production (Yi et al., 2021). In contrast, how p75^NTR^ and its downstream signaling contribute to AD tauopathy has remained unclear.

In the present study, we investigated the effects of the ΔDD and C259A signaling-deficient variants of p75^NTR^ as well as a null (knock-out) allele in the P301S mouse model of AD tauopathy. Prompted by unexpected differences between the behaviors of the ΔDD and C259A variants, we studied the effects of a mutation in the p75^NTR^ intracellular domain that specifically uncouples the receptor from the RhoA/ROCK pathway. These studies led us to discover the critical role of p75^NTR^-mediated RhoA activity in Tau-related AD pathogenesis and behavioral deficits.

## Results

### Reduced hyper-phosphorylated Tau and NFTs in hippocampus and cerebral cortex of P301S mice lacking p75^NTR^ or its death domain but not transmembrane Cys^259^

P301S mice display progressively elevated levels of hyper-phosphorylated-Tau (P-Tau) resulting from expression of the Pro^301^Ser mutant of human *MAPT* transgene. At 9 month of age, P301S mice showed comparable levels of p75^NTR^ in hippocampus (Fig. S1A-D), indicating that, unlike AD mouse models based on APP [e.g. (Yi et al., 2021)], receptor expression is not upregulated in this model of AD tauopathy. P301S mice were crossed with knock-in mice homozygous for signaling-deficient p75^NTR^ variants, i.e. either p75^ΔDD^ (lacking the dearth domain) or p75^C259A^ (Ala replacing transmembrane Cys^259^) (Tanaka et al., 2016), as well as knock-out mice lacking p75^NTR^ altogether (p75^-/-^) (Lee et al., 1992). In hippocampus of 9 month old P301S mice, p75^ΔDD^ reduced both soluble and insoluble P-Tau, the latter representing NFTs (Fig. 1A-D). Similar effects were observed in p75^-/-^ knock-out mice, although these only reached statistical significance on insoluble P-Tau (Fig. 1A-D). In contrast, p75^C259A^ had no detectable effects on P-Tau levels in hippocampus of P301S mice (Fig. 1A-D). In cerebral cortex, both p75^ΔDD^ and p75^-/-^ significantly reduced the levels of soluble and insoluble P-Tau, while again p75^C259A^ had no effect (Fig. 1E-H). Total Tau protein levels were unchanged across all genotypes (Fig. S2A, B), indicating that the observed differences in P-Tau and NFT burden resulted from altered Tau phosphorylation and aggregation rather than Tau expression.

**Figure 1.**
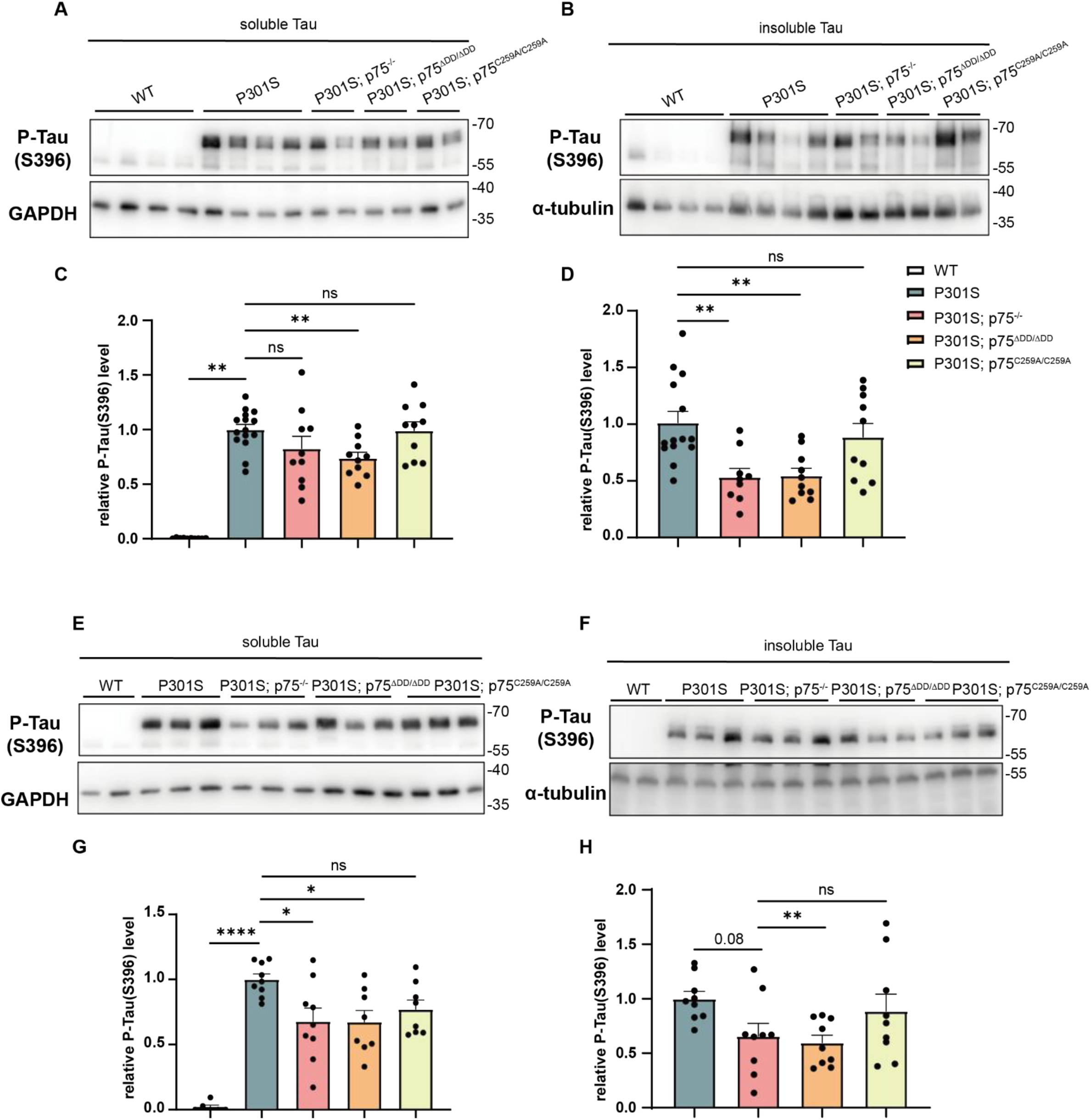
Reduced hyper-phosphorylated Tau and NFTs in hippocampus and cerebral cortex of P301S mice lacking p75^NTR^ or its death domain but not transmembrane Cys^259^. (A-B) Representative Western blots of soluble (A) and insoluble (B) P-Tau of hippocampus from 9-month-old WT, P301S, P301S/p75^-/-^, P301S/p75^ΔDD^ and P301S/p75^C259A^ as indicated using S396 antibody. Molecular weights are indicated in kDa. (C-D) Quantification of soluble (C) and insoluble (D) P-Tau from mouse hippocampus relative to GAPDH and α-tubulin, respectively. N=10 (WT), 14-15 (P301S), 9-10 (P301S/p75^-/-^), 10 (P301S/p75^ΔDD^, P301S/p75^C259A^) mice per genotype, respectively. One-way ANOVA followed by Dunnett’s multiple comparisons test, mean ± SEM. *p<0.05, **p<0.01. (E-F) Representative Western blots of soluble (A) and insoluble (B) P-Tau of cerebral cortex from 9-month-old WT, P301S, P301S/p75^-/-^, P301S/p75ΔDD and P301S/p75^C259A^ as indicated using S396 antibody. (G-H) Quantification of soluble (G) and insoluble (H) P-Tau from mouse cerebral cortex relative to GAPDH and α-tubulin, respectively. N=6 (WT), 8-9 (P301S, P301S/ p75^-/-^, P301S/p75^ΔDD^, P301S/p75^C259A^) mice per genotype, respectively. One-way ANOVA followed by Dunnett’s multiple comparisons test, mean ± SEM. *p<0.05, **p<0.01, ****p<0.0001.

### Reduced brain histopathology in hippocampus and piriform cortex of P301S mice lacking p75^NTR^ or its death domain but not transmembrane Cys^259^

Accumulation of hyper-phosphorylated Tau and NFTs is known to induce brain neuroinflammation in the form of microglial and astrocytic activation, leading to neuronal damage. Consistent with this, microgliosis and astrogliosis were significantly increased in 9-month-old hippocampus of P301S mice compared with wild type (WT), as indicated by increased immunostaining of microgliosis markers Iba1 and CD68 (Fig. 2A, B, E, F), as well as astrogliosis markers Glial fibrillary acidic protein (GFAP) and Vimentin (Fig 2C, D, G, H).

**Figure 2.**
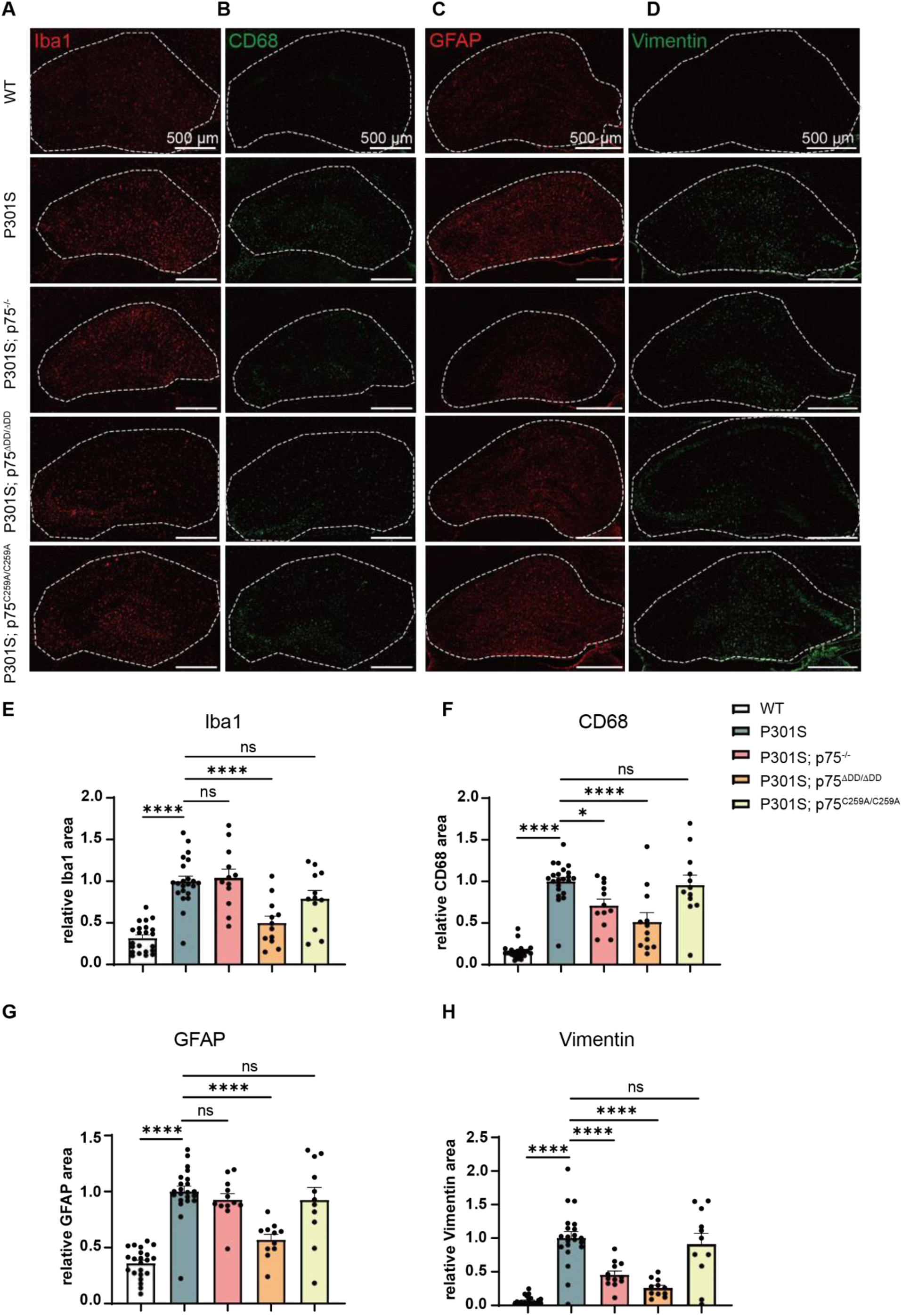
Reduced microgliosis and astrogliosis in hippocampus of P301S mice lacking p75^NTR^ or its death domain but not transmembrane Cys^259^. (A-B) Representative microgliosis confocal images of Iba1 (red) (A) and CD68 (green) (B) immunohistology in the hippocampus of 9-month-old WT, P301S, P301S/p75^-/-^, P301S/p75^ΔDD^ and P301S/p75^C259A^, as indicated. Scale bar, 500 μm. (C-D) Representative astrogliosis confocal images of GFAP (red) (C) and Vimentin (green) (D) immunohistology in the hippocampus of 9-month-old WT, P301S, P301S/p75^-/-^, P301S/p75^ΔDD^ and P301S/p75^C259A^, as indicated. Scale bar, 500 μm. (E-F) Quantification of the level of microgliosis assessed by the relative Iba1 (E) and CD68 (F) area. N=22(WT, P301S), 12(P301S/p75^-/-^, P301S/p75^ΔDD^, P301S/p75^C259A^) mice per genotype, respectively. One-way ANOVA followed by Dunnett’s multiple comparisons test, mean ± SEM. *p<0.05, ****p<0.0001. (G-H) Quantification of the level of astrogliosis assessed by the relative GFAP (G) and Vimentin (H) area. N=22(WT, P301S), 11-12(P301S/p75^-/-^, P301S/p75^ΔDD^, P301S/p75^C259A^) mice per genotype, respectively. One-way ANOVA followed by Dunnett’s multiple comparisons test, mean ± SEM. ***p<0.001, ****p<0.0001.

While Iba1 labels both resting and activated microglia in white and gray matter, CD68 marks a subset of phagocytic microglia (Fu 2014), which in the context of tauopathy is thought to reflect engagement with NFTs. In the P301S background, p75^ΔDD^ reduced hippocampal Iba1 and CD68 levels by approximately 50% (Fig. 2A, B, E, F). In p75^-/-^ mice, this only reached statistical significance in CD68^+^ microglia (Fig. 2A, B, E, F). In line with the P-Tau results, p75^C259A^ had no detectable effects on either marker (Fig. 2A, B, E, F). Analysis of the piriform cortex (PIR) of these mice revealed that both p75^ΔDD^ and p75^-/-^ could significantly reduce CD68 levels, while p75^C259A^ had again no effect (Fig. S3A, B).

GFAP expression is a widely used astrocyte marker and its expansion indicates astrocyte reactivity. On the other hand, Vimentin^+^ astrocytes are a subset of activated, disease-associated astrocytes (Chen et al., 2023; Escartin et al., 2021). In the hippocampus of P301S mice, p75^ΔDD^ significantly reduced both GFAP and Vimentin levels, while p75^-/-^ only resulted in significant amelioration of Vimentin staining, and p75^C259A^ had no effects in either (Fig. 2C, D, G, H). Analysis of PIR showed that both p75^ΔDD^ and p75^-/-^ significantly ameliorated GFAP astrogliosis, while this remained unchanged in the mice expressing p75^C259A^ (Fig. S3C, D).

### Reduced neurodegeneration and ameliorated loss of synaptic markers in P301S mice lacking p75^NTR^ death domain but not transmembrane Cys^259^

Brain degeneration is a prominent feature in advanced AD patients and aged P301S mice. To obtain a global assessment of neurodegeneration in P301S mice expressing different variants of p75^NTR^, we performed micro-computed tomography (micro-CT) to obtain a three-dimensional view of the entire mouse brain, with high contrast indicating the lateral ventricles beneath the cerebral cortex. P301S mice displayed significantly enlarged ventricle size and decreased PIR thickness compared to WT, indicating pronounced hippocampal and cortical dystrophy (Fig. 3A-C). In the P301S background, expression of p75^ΔDD^ prevented brain atrophy in both measures, with values approaching those of WT controls (Fig. 3A-C). In contrast, neither p75^-/-^ nor p75^C259A^ resulted in any significant improvement (Fig. 3A-C).

**Figure 3.**
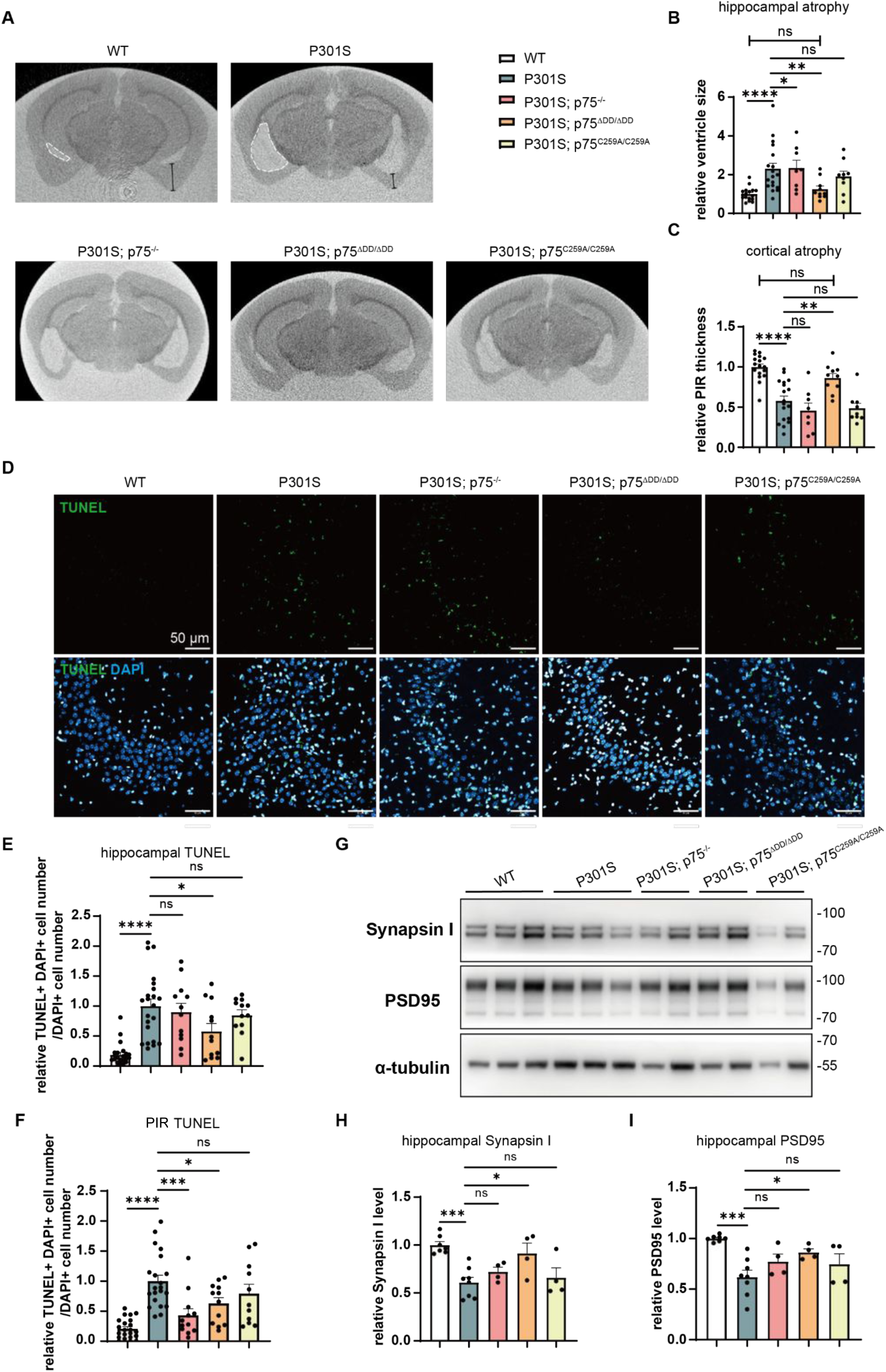
Reduced neurodegeneration and ameliorated loss of synaptic markers in P301S mice lacking p75^NTR^ death domain but not transmembrane Cys^259^. (A) Representative images from micro-CT of 10-month-old WT, P301S, P301S/p75^-/-^, P301S/p75^ΔDD^ and P301S/p75^C259A^, as indicated. White line indicates the lateral ventricle, and black line indicates the PIR thickness. (B-C) Quantification of the level of hippocampal atrophy assessed by the relative lateral ventricle size of the whole brain (B) and the relative thickness of the PIR. N=17 (WT), 18 (P301S), 8 (P301S/p75^-/-^), 10 (P301S/p75^ΔDD^), 9 (P301S/p75^C259A^) mice per genotype, respectively. One-way ANOVA followed by Holm-Sidak’s multiple comparisons test, mean ± SEM. **p<0.01, ****p<0.0001. (D) Representative confocal images of TUNEL (green) and DAPI (blue) immunohistology in hippocampal CA3 of 9-month-old WT, P301S, P301S/p75^-/-^, P301S/p75^ΔDD^ and P301S/p75^C259A^, as indicated. Scale bar, 50 μm. (E-F) Quantification of the level of dead cells assessed by the relative number of TUNEL-positive cell numbers divided by DAPI-positive cell numbers in hippocampal CA3 (E) and PIR (F). N=21-22(WT, P301S), 11-12(P301S/p75^-/-^, P301S/p75^ΔDD^, P301S/p75^C259A^) mice per genotype, respectively. One-way ANOVA followed by Dunnett’s multiple comparisons test, mean ± SEM. *p<0.05, ****p<0.0001. (G) Representative Western blots of hippocampal synaptosome extract from 9-month-old WT, P301S, P301S/p75^-/-^, P301S/p75^ΔDD^ and P301S/p75^C259A^ as indicated using PSD95 and Synapsin I antibody. Molecular weights are indicated in kDa. (H-I) Quantification of Synapsin I (H) and PSD95 (I) protein level in synaptosome from mouse hippocampus relative to α-tubulin. N=7(WT), 8(P301S), 4(P301S/p75^-/-^, P301S/p75ΔDD/ΔDD, P301S/p75^C259A^) mice per genotype, respectively. One-way ANOVA followed by Dunnett’s multiple comparisons test, mean ± SEM. *p<0.05, ***p<0.001.

Next, we assessed the extent of apoptotic cell death in hippocampus and cerebral cortex using the TUNEL assay. At 9 months of age, the number of TUNEL^+^ cells was significantly increased in P301S mice compared with WT mice (Fig. 3D), consistent with enhanced neurodegeneration driven by Tau pathology. In P301S hippocampus, TUNEL signal was significantly reduced in the p75^ΔDD^ group of mice but not in p75^-/-^ nor p75^C259A^ (Fig. 3E), in agreement with the results of micro-CT. In P301S PIR, the number of TUNEL^+^ cells was significantly reduced in both the p75^ΔDD^ and p75^-/-^ mice, while p75^C259A^ showed no effect (Fig. 3F).

Synapse loss is another hallmark of AD progression in tauopathy mouse models. To establish whether p75^NTR^ contributes to synaptic loss in P301S mice, we assessed the levels of pre- and post-synaptic proteins Synapsin I and PSD95 in synaptosome fractions isolated from the hippocampus of P301S mice carrying different p75^NTR^ alleles (Fig. 3G). The levels of both proteins were significantly decreased in synaptosome fractions from 9-month-old P301S mice expressing WT p75^NTR^, indicating substantial synaptic loss at this stage (Fig. 3H, I). Notably, analysis of P301S mice carrying different p75^NTR^ mutations revealed that p75^ΔDD^, but not p75^-/-^ nor p75^C259A^, significantly rescued the loss of Synapsin I and PSD95 in synaptosome fractions (Fig 3H, I). We note that the levels of these proteins in extracts of the whole hippocampus remained unchanged across all genotypes (Fig. S4A-C), indicating that the observed changes in synaptosome fractions reflect synaptic loss rather than altered protein expression.

Together, these results indicate that p75^NTR^ is a key determinant of neurodegeneration and synaptic loss in P301S tauopathy.

### Improved learning and memory in P301S mice lacking p75^NTR^ death domain but not transmembrane Cys^259^

Male P301S mice develop significant deficits in learning and memory by 9 months of age (Sun 2020). To evaluate the role of p75^NTR^ in spatial learning and memory in this model, we subjected P301S mice carrying different p75^NTR^ alleles to the Morris Water Maze test. P301S mice expressing WT p75^NTR^ showed significantly higher latency during the 3-day sessions of invisible platform training than WT control mice (Fig. 4A), indicating a learning deficit. We found that 9 month old P301S mice lacking p75^NTR^ (p75^-/-^) were unable to perform this test and were therefore excluded from this analysis. These mice were much thinner then those in the other groups, showed signs of self mutilation, a known feature in p75^-/-^ mice (Lee et al., 1992), and struggled to keep their head above the water surface, indicating a lack of strength. Neither p75^ΔDD^ nor p75^C259A^ P301S mice presented any of these features. Strikingly, deletion of the DD (p75^ΔDD^) almost completely reversed the learning deficits of P301S mice, with performance in the second and third training sessions that were indistinguishable from WT control mice (Fig. 4A). On the other hand, p75^C259A^ conferred no learning benefits compared to P301S mice carrying WT p75^NTR^ (Fig. 4A). We note that average swimming speed was comparable among the four groups (Fig. 4B), indicating that the differences observed in training latency were due to learning behavior rather than motor deficits.

**Figure 4.**
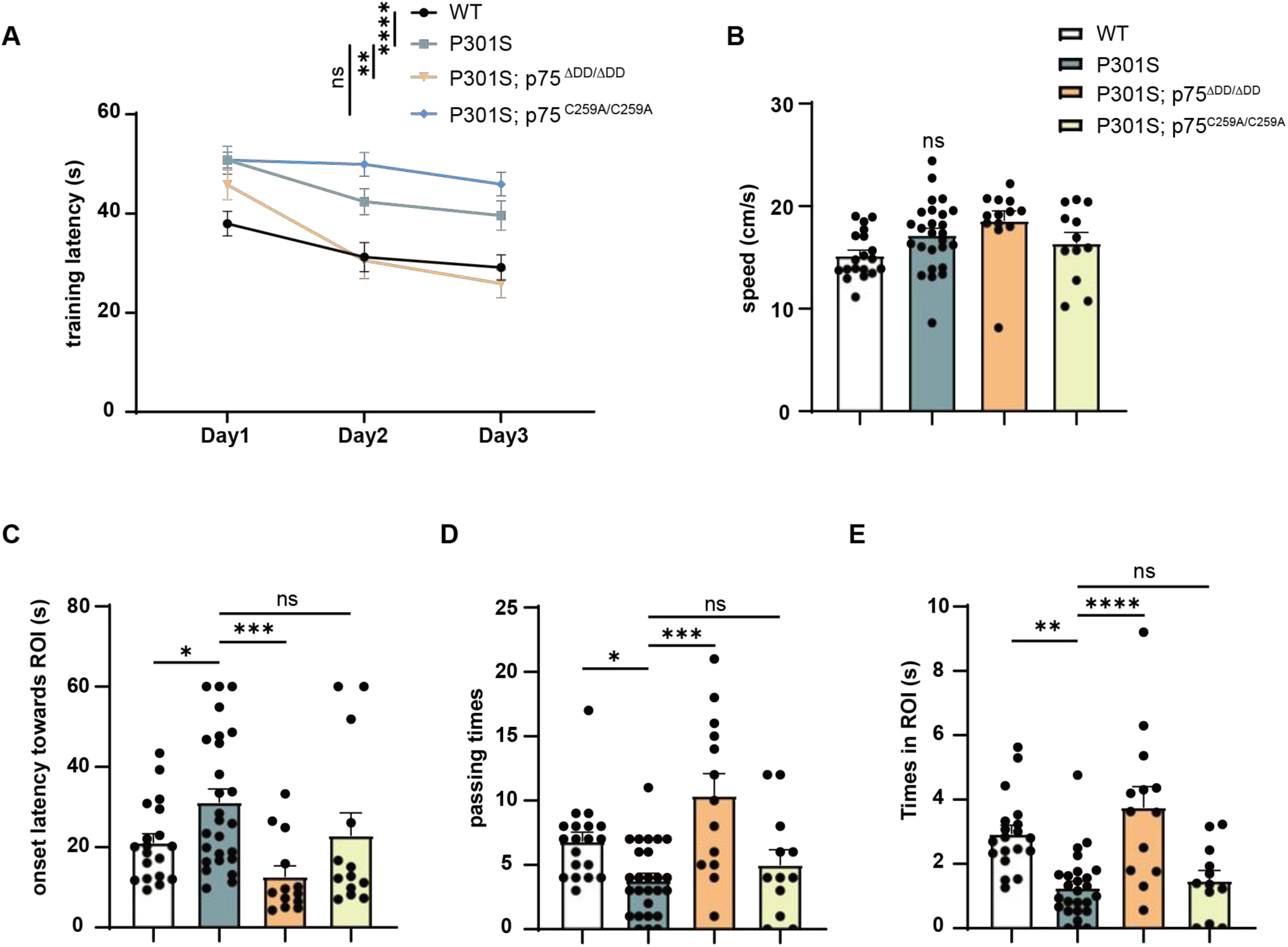
Improved learning and memory in P301S mice lacking p75^NTR^ death domain but not transmembrane Cys^259^. (A) Training latency of 3 consecutive training days of 9-month-old WT, P301S, P301S/p75ΔDD/ΔDD and P301S/p75^C259A^, as indicated. The latency of each day is an average of 5 training performed that day. N=19 (WT), 26 (P301S), 13 (P301S/p75^ΔDD^), 12 (P301S/p75^C259A^) mice per genotype, respectively. Two-way ANOVA followed by Dunnett’s multiple comparisons test, mean ± SEM. **p<0.01, ****p<0.0001. (B) Average swimming speed in the training session. One-way Ho followed by Holm-Sidak’s multiple comparisons test, mean ± SEM, ns, no significance. (C-E) Probe trial 24 h after the last training. Histograms show onset latency towards ROI (C), number of times passing ROI (D) and the time spent in ROI (E). One-way ANOVA followed by Dunn’s multiple comparisons test, mean ± SEM. *p<0.05, **p<0.01, ***p<0.001, ****p<0.0001.

A probe test (platform removed) to assess memory retention was conducted 24h after the last training session. Three parameters were analyzed in this session, namely onset latency towards the region of interest (ROI), number of times the mouse crossed over the ROI, and time spent in the ROI. P301S mice expressing WT p75^NTR^ showed significant deficits in these parameters (Fig. 4C-E). Importantly, p75^ΔDD^ rescued performance in all three parameters to nearly WT levels, while p75^C259A^ had no effect on the performance of P301S mice (Fig. 4C-E). These results indicate an important role of p75^NTR^ death domain signaling in the learning and memory deficits caused by P301S tauopathy.

### Reduced Tau pathology and behavioral deficits in P301S mice carrying RhoA-signaling deficient p75^NTR^ mutation

Our previous studies demonstrated comparable neuroprotective effects of p75^ΔDD^ and p75^C259A^ variants in neurodegeneration resulting from pilocarpine-induced seizures (Tanaka et al., 2016), as well as Aβ pathology and learning deficits in the 5xFAD mouse model of AD (Li et al., 2026b; Yi et al., 2021). Focusing on Tau-related neuropathology, the present study reveals a striking discrepancy between these two signaling-deficient p75^NTR^ variants, with p75^ΔDD^ displaying remarkable protective effects in all parameters examined, while p75^C259A^ having no effect at all. Although both variants are unable to respond to neurotrophins, one fundamental difference between them is the ability of p75^C259A^, but not p75^ΔDD^, to stimulate RhoA signaling in response to myelin-derived ligands, such as myelin-associated glycoprotein (MAG) and the Nogo peptide (Vilar et al., 2009). p75^NTR^ can enhance RhoA signaling through the recruitment of the Rho inhibitory protein RhoGDI to its DD (Yamashita and Tohyama, 2003; Lin et al., 2015). This interaction releases RhoA from RhoGDI leading to RhoA activation (Yamashita and Tohyama, 2003; Ramanujan and Ibáñez, 2024). While neurotrophins decrease RhoGDI binding to p75^NTR^, MAG, Nogo and other myelin-derived proteins increase it, and thereby potently enhance RhoA signaling through this receptor (Yamashita and Tohyama, 2003; Ramanujan and Ibáñez, 2024). Thus, although p75^ΔDD^ is totally unable to link to the RhoA pathway, p75^C259A^ still remains competent to enhance RhoA signaling in response to myelin-derived proteins (Vilar et al., 2009). We speculated that the ability of p75^C259A^ to link to the RhoA pathway may have been the reason why it failed to elicit any neuroprotection in the P301S model, prompting us to focus our attention in this pathway.

We have recently reported that the triple Ala replacement of Lys^303^, in the p75^NTR^ juxtamembrane domain, together with Lys^346^ and Glu^349^, in the p75^NTR^ DD, (herein referred to as the KKEA mutant or p75^KKEA^) completely prevents RhoGDI binding to p75^NTR^, resulting in a receptor that is no longer competent to link to the RhoA pathway (Li et al., 2026a). Importantly, and unlike p75^ΔDD^, p75^KKEA^ is still able to activate the NF-kB and JNK/caspase-3 pathways (Li et al., 2026a). In the present study, we utilized p75^KKEA^ knock-in mice (Li et al., 2026a) to examine the importance of p75^NTR^ coupling to RhoA signaling on the development of AD tauopathy in P301S mice.

In hippocampus of 9 month old P301S mice, p75^KKEA^ reduced soluble P-Tau levels but had no effect on the insoluble component (Fig. 5A-D). Regarding gliosis, the hippocampus of P301S mice expressing p75^KKEA^ showed significant reductions in Iba1, GFAP and Vimentin staining compared to P301S mice expressing WT p75^NTR^, while the reduction observed in CD68 did not reach statistical significance (Fig. 5E-H). In line with these results, micro-CT analysis showed lower lateral ventricle enlargement in p75^KKEA^ mice, although this effect did not reach statistical significance (Fig. 5I). On the other hand, TUNEL signal was significantly reduced by the KKEA mutation (Fig. 5J), indicating a lower incidence of hippocampal neuron death in P301S mice expressing p75^KKEA^. In the Morris Water Maze, test P301S mice expressing p75^KKEA^ showed significantly reduced onset latency towards ROI, indicating an amelioration effect by this mutation (Fig. 5K).

**Figure 5.**
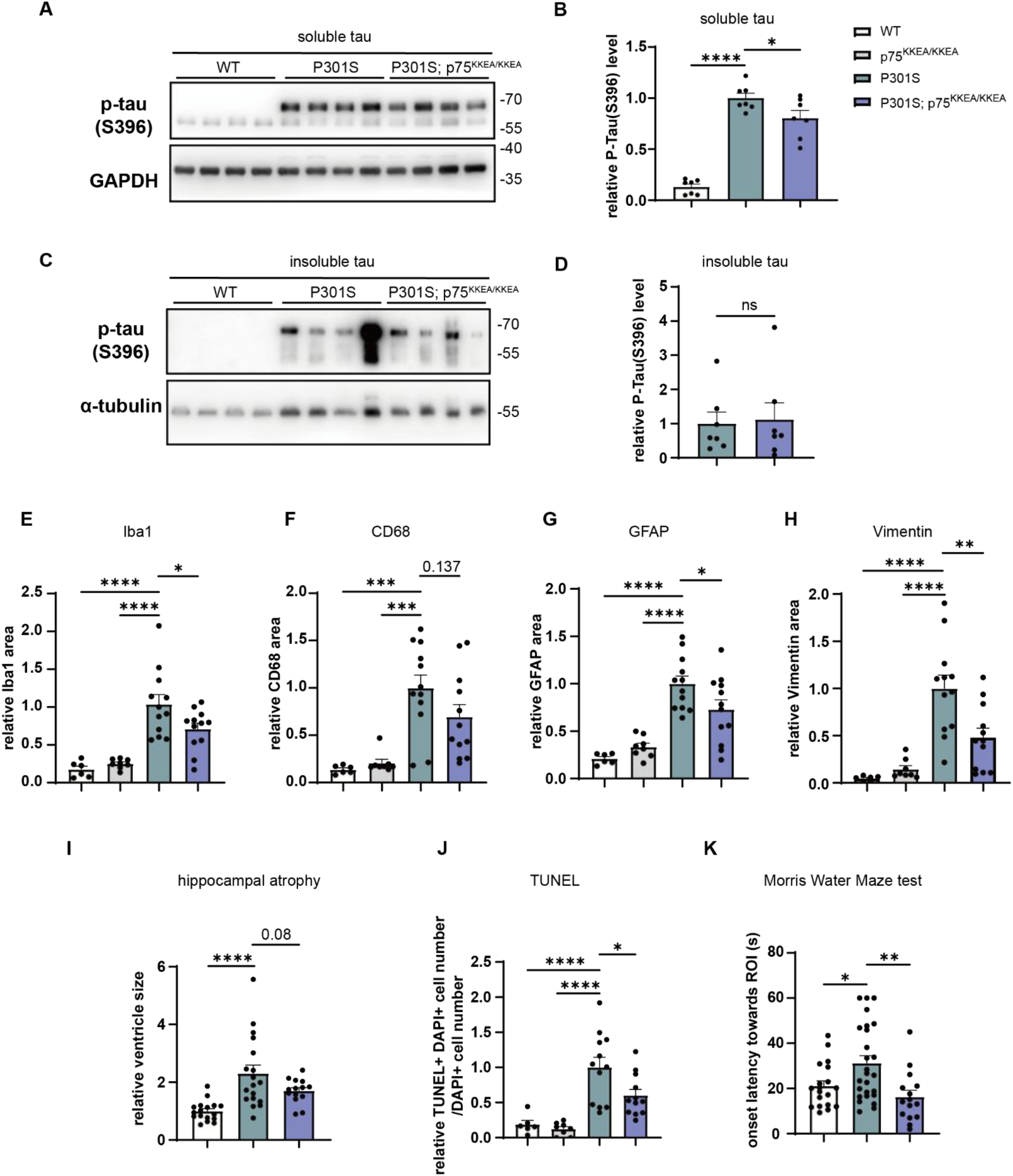
Reduced Tau pathology and behavioral deficits in P301S mice carrying RhoA-signaling deficient p75^NTR^ mutation. (A-D) Representative Western blots (A, C) and quantification results (B, D) of soluble (A, B) and insoluble (C, D) P-Tau of hippocampus from 9-month-old WT, P301S and P301S/p75^KKEA^ as indicated using S396 antibody relative to GAPDH and α-tubulin, respectively. Molecular weights are indicated in kDa. N=7 mice per genotype. One-way ANOVA followed by Dunnett’s multiple comparisons test, mean ± SEM. *p<0.05, ****p<0.0001. (E-H) Quantification of the level of microgliosis and astrogliosis assessed by the relative Iba1 (E), CD68 (F), GFAP (G) and Vimentin (H) area in the hippocampus of 9-month-old WT, p75^KKEA^, P301S and P301S/p75^KKEA^, as indicated. N=6 (WT), 8 (p75^KKEA^), 12 (P301S, P301S/p75^KKEA^) mice per genotype, respectively. One-way ANOVA followed by Dunnett’s multiple comparisons test, mean ± SEM. *p<0.05, **p<0.01, ***p<0.001, ****p<0.0001. (I) Quantification of the level of hippocampal dystrophy assessed by the relative lateral ventricle size of the whole brain of 10-month-old WT, P301S and P301S/p75^KKEA^, as indicated. WT and P301S are the same as in Fig. 3B. N=17(WT), 18(P301S), 14(P301Sp75^KKEA^) mice per genotype, respectively. One-way ANOVA followed by Dunnett’s multiple comparisons test, mean ± SEM. ****p<0.0001. (J) Quantification of the level of dead cells assessed by the relative number of TUNEL-positive cell numbers divided by DAPI-positive cell numbers in the hippocampal CA3 of 9-month-old WT, p75^KKEA^, P301S and P301S;p75^KKEA^, as indicated. N=6 (WT), 8 (p75^KKEA^), 12 (P301S, P301S/p75^KKEA^) mice per genotype, respectively. One-way ANOVA followed by Dunnett’s multiple comparisons test, mean ± SEM. *p<0.05, ****p<0.0001. (K) Quantification of the probe trial of the Morris Water Maze test 24 h after the last training. Histograms show onset latency towards ROI of 9-month-old WT, P301S and P301S/p75^KKEA^, as indicated. WT and P301S are the same as in Fig. 4C. N=19 (WT), 26 (P301S), 14 (P301S/p75^KKEA^) mice per genotype, respectively. One-way ANOVA followed by Dunnett’s multiple comparisons test, mean ± SEM. *p<0.05, **p<0.01.

Together, these results demonstrate a protective effect of p75^KKEA^ in P301S tauopathy, indicating a critical role of p75^NTR^ signaling to the RhoA pathway in the pathophysiology of AD tauopathy.

### p75^NTR^-mediated ROCK activity is critical for the hyper-phosphorylation of Tau protein in P301S neurons

To further dissect the mechanisms underlying the effects of p75^NTR^-mediated RhoA signaling in AD Tau pathology we utilized primary neuron cultures derived from P301S mice (or infected with AAV-DJ-P301S virus) expressing different p75^NTR^ variants (Fig. S5). In the first set of experiments, we confirmed the role of RhoA/ROCK signaling in Tau hyper-phosphorylation in cultures of cortical neurons using a battery of pharmacological activators and inhibitors. Treatment with the RhoA activator lysophosphatidic acid (LPA) increased ROCK activity in cortical neurons from P301S mice (Fig. S6) and elevated Tau phosphorylation in WT cortical neurons infected with AAV-DJ-P301S (Fig S7A, B). Conversely, treatment with the RhoA inhibitor Rhosin, or the ROCK inhibitor Fasudil, suppressed ROCK activity (Fig. S6) and reduced Tau phosphorylation (Fig. S7C-F). Finally, inhibition of GSK3β, the major Tau kinase downstream of ROCK (Medd et al., 2025), with Laduviglusib significantly reduced P-Tau levels in P301S neurons cultures (Fig. S7G, H). These results confirmed that P-Tau levels in cortical neurons expressing Tau^P301S^ is under the control of RhoA/ROCK signaling.

Consistent with our results from tissue extracts, P301S cortical neurons lacking p75^NTR^ (p75^-/-^) or expressing p75^ΔDD^ or p75^KKEA^ showed reduced ROCK activity and Tau phosphorylation levels compared to P301S neurons expressing WT p75^NTR^ (Fig.6A-I). As expected, ROCK activity and P-Tau levels remained unchanged in P301S neurons expressing p75^C259A^ (Fig. 6J-L). Similar results were obtained in P301S hippocampal neurons (Fig. S8A-H). The level of activated GSK3β was decreased in p75^-/-^, p75^ΔDD^ and p75^KKEA^ P301S cortical neurons, as reflected by increased phosphorylated GSK3β relative to total GSK3β, while remaining unchanged in P301S neurons expressing p75^C259A^ (Fig. S9A-J). We note that activated GSK3β levels were unaffected by the P301S transgene itself (Fig. S9A-B). Finally, stimulation with MAG increased ROCK activity in cortical neurons expressing wild type p75^NTR^ and p75^C259A^, but not in P301S neurons lacking p75^NTR^ (p75^-/-^) or expressing p75^ΔDD^ or p75^KKEA^ (Fig. 7A-E). In agreement with this, MAG treatment enhanced P-Tau levels in AAV-DJ-P301S-infected cortical neurons derived from WT and p75^C259A^ mice, but not from p75^-/-^, p75^ΔDD^ or p75^KKEA^ mice (Fig. 7F-O).

**Figure 6.**
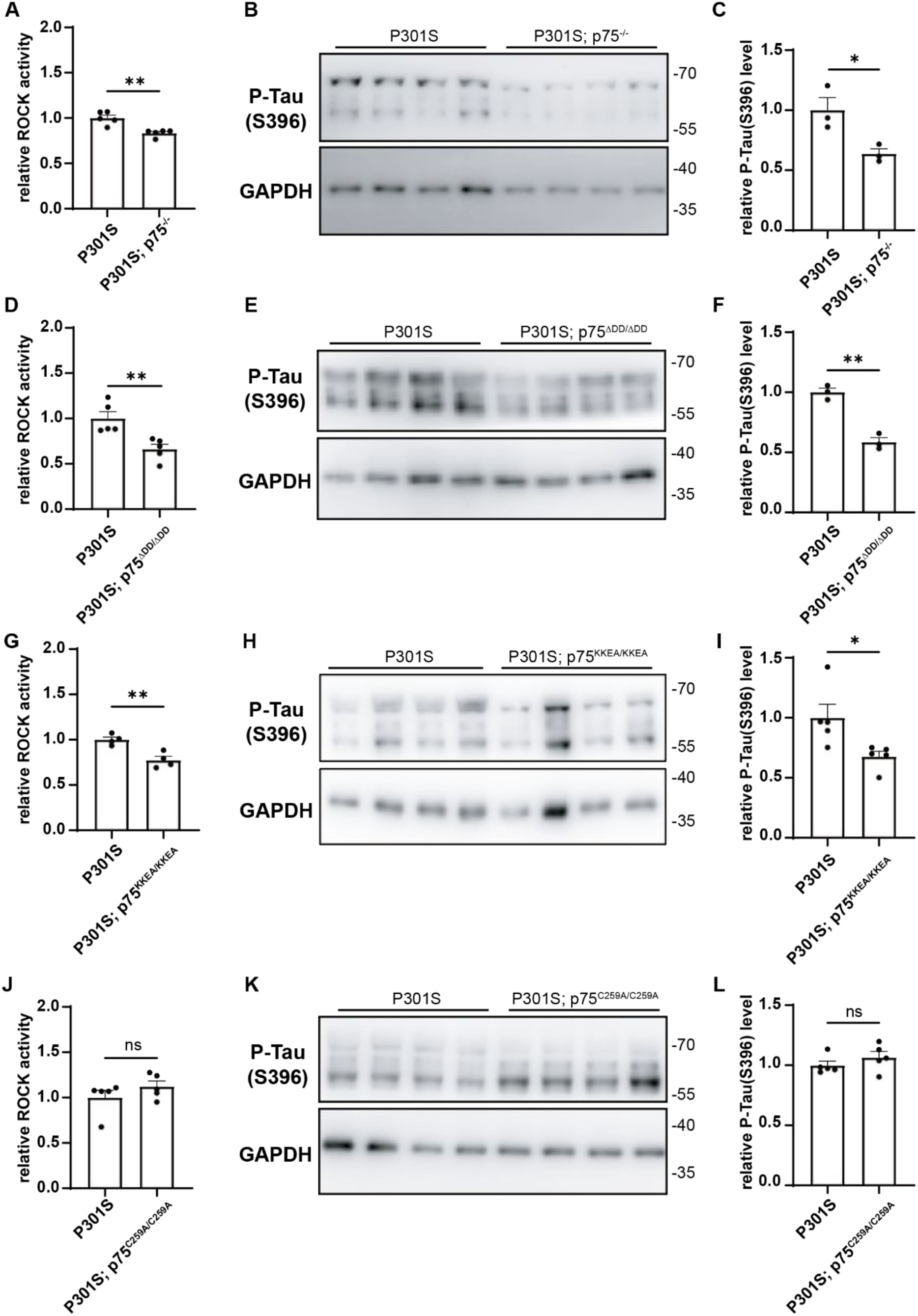
Reduced P-Tau levels and ROCK activity in cultured cerebral cortical neurons of P301S mice carrying KO, ΔDD and KKEA but not C259A p75^NTR^ variants. (A) Quantification results of ROCK ELISA from P301S and P301S/p75^-/-^ neurons. Results shown are from 5 independent experiments with 2 replicates each. Unpaired Student’s t-test, mean ± SEM. **p<0.01. (B) Representative Western blots of cortical neurons from P301S and P301S/p75^-/-^ embryos, as indicated. Molecular weights are indicated in kDa. (C) Quantification results of Western blot from P301S and P301S/p75^-/-^ neurons. Results shown are from 3 independent experiments with 4 replicates each. Unpaired Student’s t-test, mean ± SEM. *p<0.05. (D) Quantification results of ROCK ELISA from P301S and P301S/p75^ΔDD^ neurons. Results shown are from 5 independent experiments with 2 replicates each. Unpaired Student’s t-test, mean ± SEM. **p<0.01. (E) Representative Western blots of cortical neurons from P301S and P301S/p75^ΔDD^ embryos, as indicated. (F) Quantification results of Western blot from P301S and P301S/p75^ΔDD^ neurons. Results shown are from 3 independent experiments with 4 replicates each. Unpaired Student’s t-test, mean ± SEM. **p<0.01. (G) Quantification results of ROCK ELISA from P301S and P301S/p75^KKEA^ neurons. Results shown are from 4 independent experiments with 2 replicates each. Unpaired Student’s t-test, mean ± SEM. **p<0.01. (H) Representative Western blots of cortical neurons from P301S and P301S/p75^KKEA^ embryos, as indicated. (I) Quantification results of Western blot from P301S and P301S/p75^KKEA^ neurons. Results shown are from 5 independent experiments with 4 replicates each. Unpaired Student’s t-test, mean ± SEM. *p<0.05. (J) Quantification results of ROCK ELISA from P301S and P301S/p75C^259A^ neurons. Results shown are from 5 independent experiments with 2 replicates each. Unpaired Student’s t-test, mean ± SEM. No significant difference between P301S/p75^C259A^ versus P301S. (K) Representative Western blots of cortical neurons from P301S and P301S/p75C^259A^ embryos, as indicated. (L) Quantification results of Western blot from P301S and P301S/p75C^259A^ neurons. Results shown are from 5 independent experiments with 4 replicates each. Unpaired Student’s t-test, mean ± SEM, ns, no significance.

**Figure 7.**
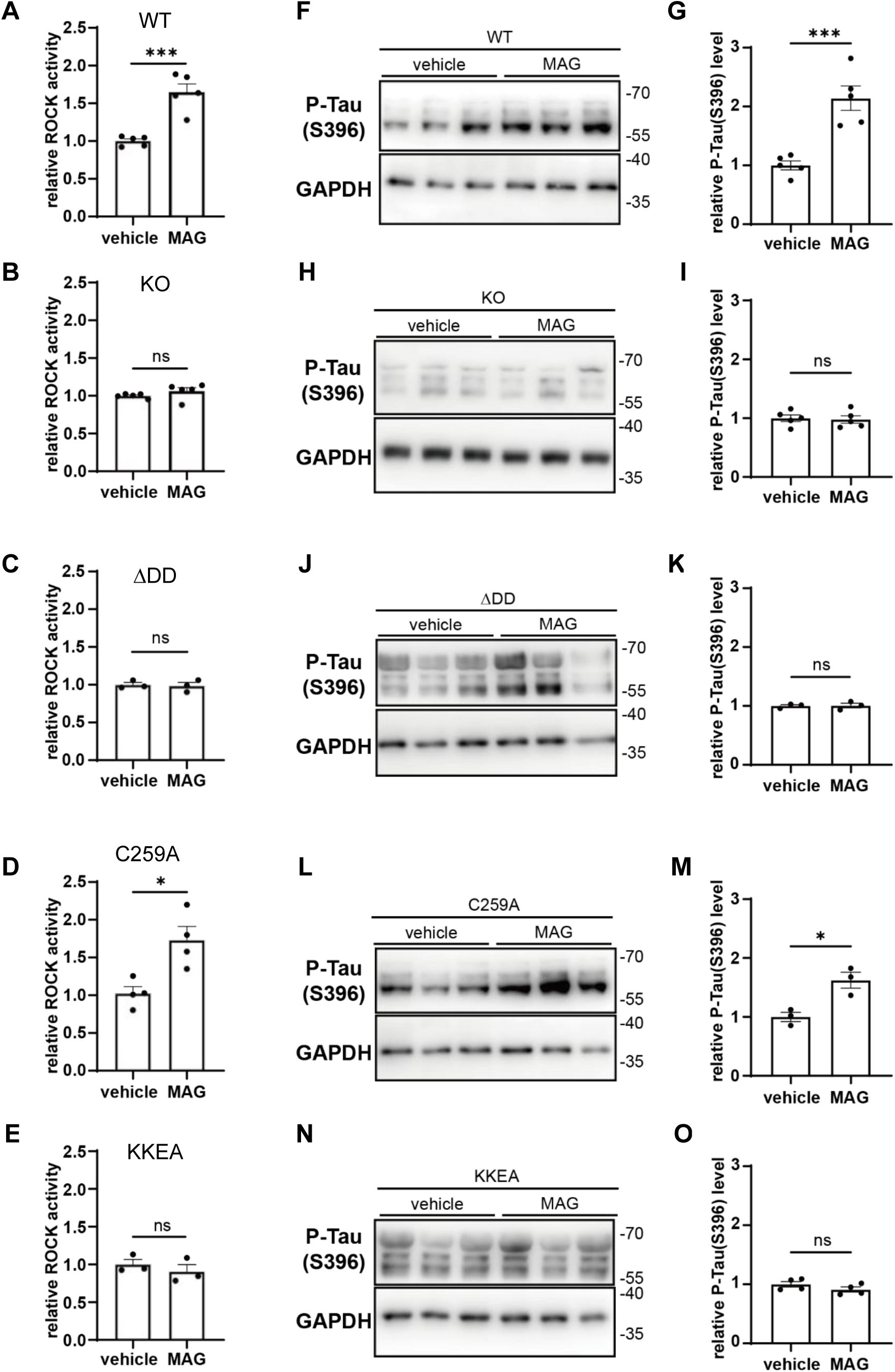
ROCK activity and P-Tau levels after MAG stimulation of wild type, p75^-/-^, p75^ΔDD^, p75C^259A^ and p75^KKEA^ cortical neurons. (A-E) Quantification of ROCK activity in WT (A), p75^-/-^ (B), p75^ΔDD^ (C), p75C^259A^ (D) and p75^KKEA^ (E) cortical neurons. Results shown are from 3 to 5 independent experiments with 2 replicates each. Unpaired Student’s t-test, mean ± SEM. *, p<0.05/***, p<0.001/ns, no significance. (F, G) Representative Western blots (F) and P-Tau quantification normalized to GAPDH levels (G) of cortical neurons derived from WT embryos infected with AAV-DJ-P301S virus and treated with either vehicle or MAG (2μg/mL) for 24h, as indicated. Molecular weights are indicated in kDa. Results shown are from 5 independent experiments with 3 replicates each. Unpaired Student’s t-test, mean ± SEM/***, p<0.001. (H, I) Representative Western blots (H) and P-Tau quantification normalized to GAPDH levels (I) of cortical neurons derived from p75^-/-^ embryos infected with AAV-DJ-P301S virus and treated with either vehicle or MAG (2μg/mL) for 24h, as indicated. Molecular weights are indicated in kDa. Results shown are from 4 independent experiments with 3 replicates each. Unpaired Student’s t-test, mean ± SEM; ns, no significance. (J, K) Representative Western blots (K) and P-Tau quantification normalized to GAPDH levels (K) of cortical neurons derived from p75^ΔDD^ embryos infected with AAV-DJ-P301S virus and treated with either vehicle or MAG (2μg/mL) for 24h, as indicated. Molecular weights are indicated in kDa. Results shown are from 3 independent experiments with 3 replicates each. Unpaired Student’s t-test, mean ± SEM, ns, no significance. (L, M) Representative Western blots (L) and P-Tau quantification normalized to GAPDH levels (M) of cortical neurons derived from p75C^259A^ embryos infected with AAV-DJ-P301S virus and treated with either vehicle or MAG (2μg/mL) for 24h, as indicated. Molecular weights are indicated in kDa. Results shown are from 3 independent experiments with 3 replicates each. Unpaired Student’s t-test, mean ± SEM. *, p<0.05. (N, O) Representative Western blots (N) and P-Tau quantification normalized to GAPDH levels (O) of cortical neurons derived from p75^KKEA^ embryos infected with AAV-DJ-P301S virus and treated with either vehicle or MAG (2μg/mL) for 24h, as indicated. Molecular weights are indicated in kDa. Results shown are from 4 independent experiments with 3 replicates each. Unpaired Student’s t-test, mean ± SEM, ns, no significance.

Together, these results support the notion that p75^NTR^ contributes to AD tauopathy by regulating the activation of the RhoA-ROCK pathway.

## Discussion

Of the 182 randomized clinical trials for AD ongoing in 2024, one third targeted β-amyloid and Tau, and of these, only one combined anti-β amyloid and anti-tau monoclonal antibodies (Cummings et al., 2025). The majority of ongoing trials targeted a range of other mechanisms, including the gut–brain axis, vasculature, epigenetics, circadian rhythm, growth factors, APOE status, lipid metabolism, neurogenesis, oxidative stress, protein metabolism, bioenergetics, synaptic plasticity, neurotransmitter receptors, inflammation, and immunity. This variety reflects a shift towards more complex and explanatory pathophysiological models that accommodate co-pathology and resilience (Frisoni et al., 2025). p75^NTR^ affects many of the above mentioned processes, including β-amyloid and Tau NFTs, making it a potential one-stop target for AD therapy. Encouragingly, a recent Phase 2a clinical trial of a small molecule presented as a “modulator” of p75^NTR^ activity resulted in measurable improvements in several AD biomarkers as well as mild amelioration of cognitive functions, although the latter did not reach statistical significance (Shanks et al., 2024). It is currently unknown how this drug affects p75^NTR^ signaling. As p75^NTR^ plays both positive and negative roles in different brain cell types during AD (see e.g. (Han et al., 2025)), a better understanding of the distinct p75^NTR^ mechanisms that contribute to AD pathogenesis may allow the development of more effective strategies targeting this receptor.

Two previous studies assessed the effects of a null mutant of p75^NTR^ on the P301L mouse model of tauopathy and found reduced Tau hyper-phosphorylation but no cognitive improvement at 6 months of age (Shen et al., 2018; Mañucat-Tan et al., 2019). Compared to P301L, the P301S model is more severe and has a more aggressive disease course with earlier onset, pronounced neuronal death and increased mortality (Yoshiyama et al., 2007). In our studies, we found that 9 month old P301S mice that were knock-out for p75^NTR^ were too frail to undergo the Morris Water Maze test. However, both deletion of the p75^NTR^ death domain or the KKEA mutation that uncouples RhoA signaling resulted in significant improvement in the performance of P301S mice in this test. Together with our previous study examining the effects of p75^ΔDD^ and p75^C259A^ in the 5xFAD model of β-amyloid pathology, these studies suggest that it is more beneficial to cripple receptor signaling than to remove p75^NTR^ altogether. Interestingly, one thing the p75^ΔDD^, p75^C259A^ and p75^KKEA^ variants have in common is the preservation of the receptor extracellular domain, suggesting that this domain may have neuroprotective functions, as it has been proposed in two earlier studies (Yao et al., 2015; Wang et al., 2016).

This is the first study where we find a significant difference between the p75^ΔDD^ and p75^C259A^ variants in the context of neurodegeneration. In our previous studies, both variants conferred comparable protection against neuronal death induced by pro-neurotrophins and β-amyloid (Tanaka et al., 2016; Yi et al., 2021), neurodegeneration following pilocarpine-induced seizures (Tanaka et al., 2016), and Aβ accumulation and cognitive deficits in 5xFAD mice (Yi et al., 2021; Li et al., 2026b). Unlike, p75^ΔDD^, p75^C259A^ retains the ability to stimulate RhoA signaling in response to myelin-derived ligands, such as MAG (this study and (Vilar et al., 2009)), which indicated that p75^NTR^ coupling this particular pathway may play a significant role in Tau pathology. Indeed, taking advantage of of the p75^KKEA^ variant that uncouples the receptor from RhoGDI and RhoA signaling, we found that this mutation ameliorated all disease markers in the P301S model, including Tau hyper-phosphorylation, micro- and astro-gliosis, hippocampal atrophy, neuronal cell death and spatial memory. This result suggest that uncoupling p75^NTR^ from RhoGDI/RhoA signaling may have beneficial effects in AD. This could potentially be achieved by small molecules that tweak the activity of p75^NTR^ in a selective way, perhaps by targeting the transmembrane domain of the receptor, as shown in previous work from our laboratory (Goh et al., 2018; Lopes-Rodrigues et al., 2025). In this context, it would be of interest to learn whether the compound used in the clinical trials by Shank et al. (2024) affects this pathway. Notwithstanding the effects of p75^KKEA^, we would like to note that p75^ΔDD^ was more effective than p75^KKEA^ at ameliorating Tau pathology in P301S mice, suggesting that p75^NTR^ is likely to contribute to Tau-related pathogenesis through additional pathways, perhaps those related to the activation of JNK, which p75^KKEA^ is still capable of doing. In conclusion, we find that p75^NTR^ contributes to AD tauopathy in great measure by enhancing the activity of the RhoA-ROCK pathway. Our results also reinforce the notion that impairing p75^NTR^ signaling is more beneficial in neurodegeneration than blunting receptor expression. It would be interesting to ascertain whether p75^KKEA^ is also neuroprotective in the context of β-amyloid pathology. Were this to be the case, it would make a very strong rationale for specifically targeting p75^NTR^ coupling to RhoGDI/RhoA signaling as a novel approach to the treatment of AD.

## Materials and Methods

### Mice

Mice had access to normal chow diet and water ad libitum and were maintained on a 12 h light/dark cycle. The mouse lines utilized in this study have been described previously and are as follows: P301S (Allen 2002), exon 3 knockout p75^-/-^ (Lee et al., 1992); p75^ΔDD^ and p75^C259A^ (Tanaka et al., 2016) and p75^KKEA^ (Li et al., 2026a). All mice were of C57BL/6J background. P301S mice develop significant deficits in learning and memory by 9 months of age, with reported sex dimorphism in disease severity (Sun et al., 2020). In this study, we used male mice, which exhibit more severe pathology (Sun et al., 2020). Animal care and experimental procedures were approved by Laboratory Animal Welfare and Ethics Committee of Chinese Institute for Brain Research (CIBR-IACUC-028).

### Tau protein fractionation and synaptosome extraction

Dissected frozen tissue was homogenized using RIPA Buffer (Sigma R0278) and minced, and kept on ice for 30 min, then centrifuged at 15000 rpm for 15 min at 4 °C. The supernatant was transferred and kept in aliquots as the soluble fraction, and protein concentration was measured using a BCA kit (Solarbio PC0200). The pellet was lysed by adding 4x Laemmli reagent (Bio-Rad 1610747) directly and boiled at 95 °C for 15 min, after which H2O was used to dilute to 1x solution, and this was used as the insoluble fraction.

Dissected frozen hippocampal tissue was weighted and added the Syn-PER Reagent (Thermo 87793) and minced gently. Homogenate was centrifuged at 1200 g for 10 min at 4 °C, and the supernatant was transferred to a new tube (whole tissue extract). Centrifuged the supernatant at 15000 g for 20 min at 4 °C and removed the supernatant (cytosolic fraction). Use Syn-PER Reagent to resuspend the pellet (synaptosome fraction). We used whole tissue extract to measure the protein concentration.

### Western blotting

10% SDS-PAGE gel (Bio-Rad) was used for electrophoresis (Bio-Rad). Proteins were transferred to PVDF membranes (Bio-Rad) using a semi-dry transfer system (Bio-Rad) and blocked using 5% skim milk (Solarbio D8340) for 1 h. The blots were incubated with primary antibodies at 4 °C overnight and with secondary antibodies (CST) at room temperature (RT) for 1 h. The blots were developed using Chemiluminescent HRP substrate (Millipore WBKLS0500) or Sensitivity chemiluminescent substrate (Thermo A38555) and exposed using AMERSHAM ImageQuant 800. The analysis and statistics were performed by using ImageJ software (NIH). Primary antibodies included P-Tau S396 (CST 9632), total Tau D1M9X (CST 46687), p75^NTR^ ECD (R&D AF1157), Synapsin I (Millipore AB1543), PSD95 (Sysy N3783), p-GSK3β (CST 9323), total GSK3β (CST 9315), GAPDH (Sigma-Aldrich G9545) and α-tubulin (Proteintech11224-1-AP).

### Immunohistochemistry and image analysis

Cryostat sections (14 μm) of mouse brain were washed with PBS and pre-blocked in blocking buffer with PBS containing 5% normal donkey serum (Jackson Immuno 017-000-121) and 0.2% Triton X-100 for 30 min at RT. Then sections were incubated with primary antibodies at 4 °C overnight and with secondary antibodies (Invitrogen) at RT for 1 h. Before mounting with mounting medium (Dako S3023), sections were stained with DAPI (Solarbio C0065) for 10 min and washed with PBS. The imaging was performed using a confocal microscope (Zeiss LSM900) and the analysis and statistics were performed by ImageJ software. Primary antibodies included p75NTR ECD, Iba1(Wako 019-19741), CD68 (Bio-Rad MCA1957), GFAP (Abcam ab7260) and Vimentin (Thermo PA1-10003).

TUNEL was performed using DeadEndTM Fluorometric TUNEL System (Promega G3250). Briefly, sections were permeabilized with Proteinase K solution for 5 min at RT, and Equilibration Buffer was added for 10 min at RT. Then the sections were incubated with TdT reaction mix for 1 h at 37 °C, avoid from light. 2x SSC was added to the sections to stop reaction for 15 min at RT, followed by PBS wash and mounting.

For each mouse, three to five brain coronal sections were quantified. For p75^NTR^ and gliosis markers, positive signal area was normalized to total area of hippocampus. For quantification of TUNEL signal, stacks of 18 consecutive confocal images taken at 0.5 μm intervals were acquired and maximum projection images were used for quantification. The TUNEL and DAPI double positive cell numbers divided by total cell numbers was used as results for TUNEL.

### Micro CT

Mice were perfused and the brains were post-fixed with 4% paraformaldehyde at 4 °C for 24 h, after which the brains would be merged in 35% iohexol (Aladdin I134719) for 2 weeks at 4 ℃. Micro-CT scanning was done by NEMO Micro-CT (NMC-200) and the Cruser software. The chamber used was the ex vivo large scanning chamber. The analysis was done using Avatar3 software. The CT reconstruction method was FDK and resolution was 1K*1K. The ventricle volume quantification included brain region starting from around Bregma -1.82 mm and ending at where the ventricles disappeared.

### Morris Water Maze test

The mice were acclimated to a reversed light-dark cycle before the test. The training includes visible-platform (1 day) and invisible-platform (3 days) phase. Each mouse underwent five trials per training day with 1 min per training, and a circular platform remained in a fixed position starting from the first day of invisible-platform phase for mice to stand on and rest. 24 h after the last training, a probe trial was performed by removing the platform and recording the swimming for 1 min. Swimming path and time were recorded and analyzed. During the experiment, mice were carefully dried and placed near a heating source to prevent hypothermia after each trial.

### Primary culture, AAV virus and ROCK ELISA

Primary hippocampal and cerebral cortical neurons were cultured from embryonic 17.5 mice. The tissues were dissected in dissection buffer (HEPES (Gibco 15630-080), Leibovitz’s-15 medium (Gibco 21083-027)) and were subjected to pre-warmed digestion buffer (papain (Worthington LS003126) and DNase I (Roche 10104159001) in dissection buffer) for a 30 min incubation at 37 °C. Then the solution was centrifuged at 1000 rpm for 3 min, and pellet was resuspended with feeding media (HEPES, GlutaMax (Gibco 35050-061), B27 supplement (Thermo 17504-044) in Neurobasal medium (Gibco 21103-049)). The tissues were triturated by gentle pipetting and went through 70 μm filter membrane, followed by a 3 min centrifugation at 100 g. Lastly, the neurons were resuspended in feeding media and seed to plates pre-coated with PDL (Sigma-Aldrich P7405) and Laminin (Sigma-Aldrich L2020). The primary neurons were cultured in a 37 °C incubator of 5% CO2.

AAV-DJ-GFP and -P301S virus were purchased from Vigene Biosciences. The viral vector was pAV-hsyn. Virus transduction of primary cortical neurons was performed after 3 days in vitro (DIV3) at multiplicity of infection (MOI) = 105 and left for 5 days before addition of drugs.

ROCK ELISA was performed using ROCK Activity Assay Kit (Abcam ab211175). Briefly, primary cerebral cortical neurons were harvest at DIV5 with lysis buffer and centrifuged at 15000 rpm for 1 min at 4 °C. Supernatant was transferred and added to ELISA plate, and Kinase Reaction Buffer/DTT/ATP mix was added and mixed by pipetting, followed by 1 h incubation at 30 °C with gentle agitation. Then primary p-MYPT1 antibody and HRP-conjugated secondary antibody were added and incubated for 1 h at RT separately, followed by substrate solution incubation for 15 min at RT. Added stop solution right before reading the plate using a multimode microplate reader (BioTek) at 450 nm.

### Statistical analysis

Statistics analyses were performed using Prism software. Results are presented as mean ± SEM. Student’s t-test, one-way ANOVA or two-way ANOVA was performed to test statistical significance according to the requirements of the experiments. One-way ANOVA followed by Dunnett’s multiple comparisons test or Holm-Sidak’s multiple comparisons test was used. Behavioral data were analyzed using two-way ANOVA followed by Dunnett’s multiple comparisons test. Statistical significance: *p<0.05, **p<0.01, ***p<0.001, ****p<0.0001.

## Acknowledgements

The authors would like to thank Lei Wang, Jocelyn Jia, Yankui Fu and Shuo Zhang for technical and admin assistance. This work was supported by research grants to C.F.I. from Peking University, Chinese Institute for Brain Research, Beijing, and Swedish Research Council (Vetenskapsrådet, contract nr. 2024-03222); and a startup grant to M.X. from Swedish Research Council (Vetenskapsrådet, contract nr. 2021-01805).

## Author contributions

K.L. performed all experimental work, analyzed data and prepared a draft of the manuscript and figures; M.X. co-directed the project and corrected the manuscript; C.F.I. conceived the project, directed the research and wrote the final version of the manuscript.

## Supplementary Figures

**Figure S1.**
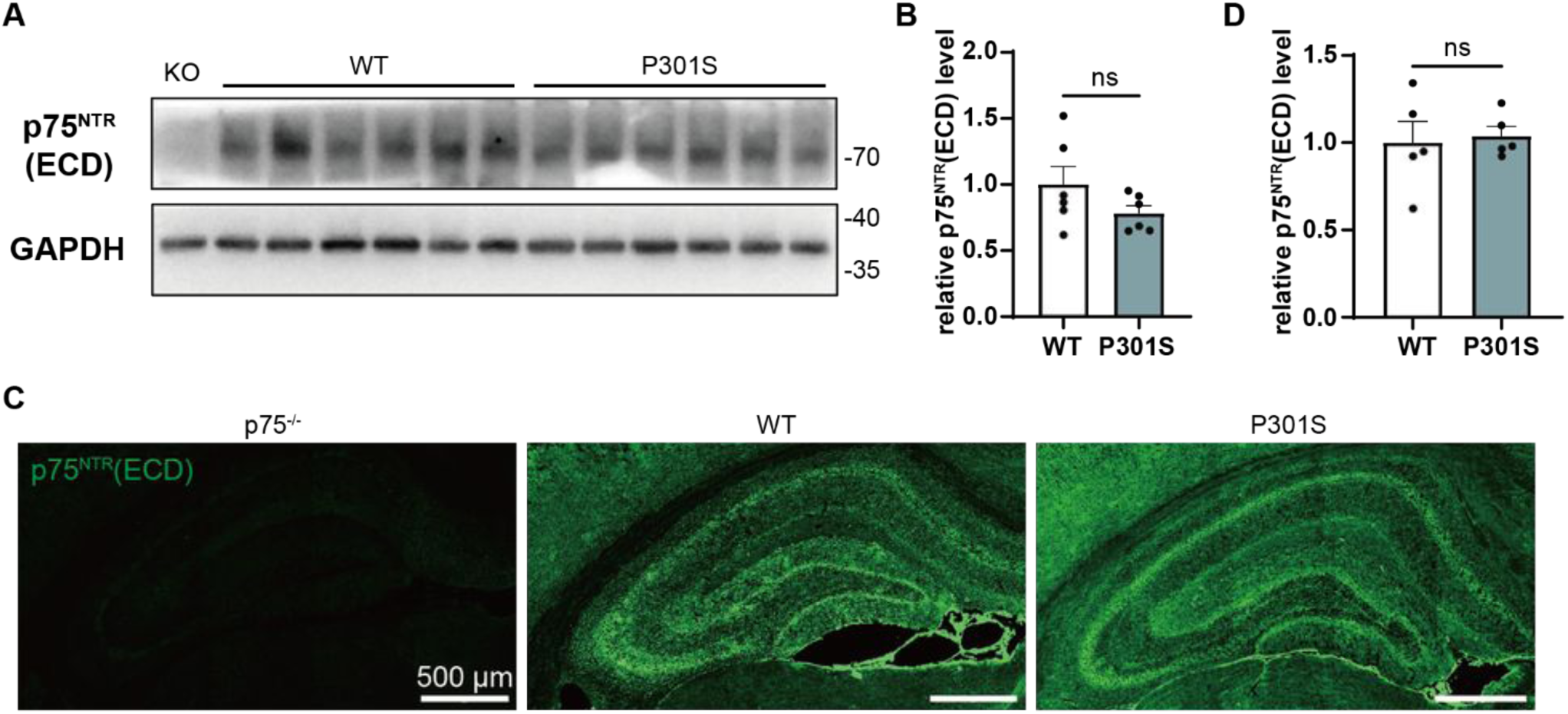
Comparable p75^NTR^ level between P301S and WT. A) Representative Western blot gels from hippocampus of 9-month-old p75^-/-^, WT and P301S as indicated using p75^NTR^ (ECD) antibody. Molecular weights are indicated in kDa. B) Quantification of p75^NTR^ from hippocampus relative to GAPDH. N=6 mice per genotype. Unpaired Student’s t-test, mean ± SEM. No significant difference between WT versus P301S. C) Representative confocal images of p75^NTR^ (green) immunohistology in the hippocampus of 9-month-old p75^-/-^, WT and P301S, as indicated. Scale bar, 500 μm. D) Quantification of the level of p75^NTR^. N=5 mice per genotype. Unpaired Student’s t-test, mean ± SEM. No significant difference between WT versus P301S.

**Figure S2.**
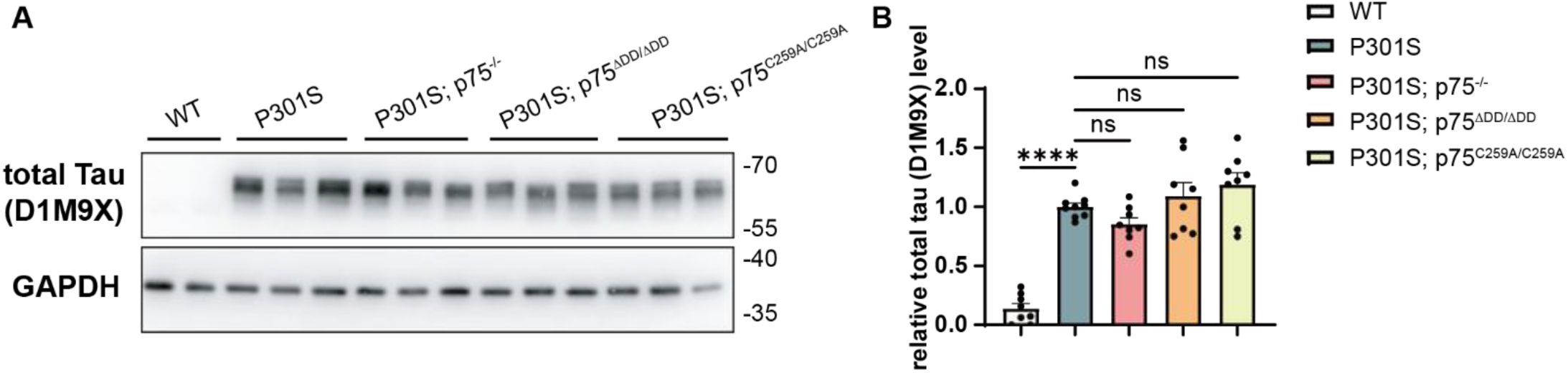
Comparable total Tau expression level among WT and transgenic mouse models. A) Representative Western blot gels from hippocampus of 9-month-old WT, P301S, P301S; p75^-/-^, P301S; p75^ΔDD/ΔDD^ and P301S; p75^C259A/C259A^ as indicated using D1M9X antibody. Molecular weights are indicated in kDa. B) Quantification of total Tau from hippocampus relative to GAPDH. N=8 mice per genotype, respectively. One-way ANOVA followed by Dunnett’s multiple comparisons test, mean ± SEM. ****p<0.0001, no significant difference between P301S; p75^-/-^, P301S; p75^ΔDD/ΔDD^, P301S; p75^C259A/C259A^ versus P301S, respectively.

**Figure S3.**
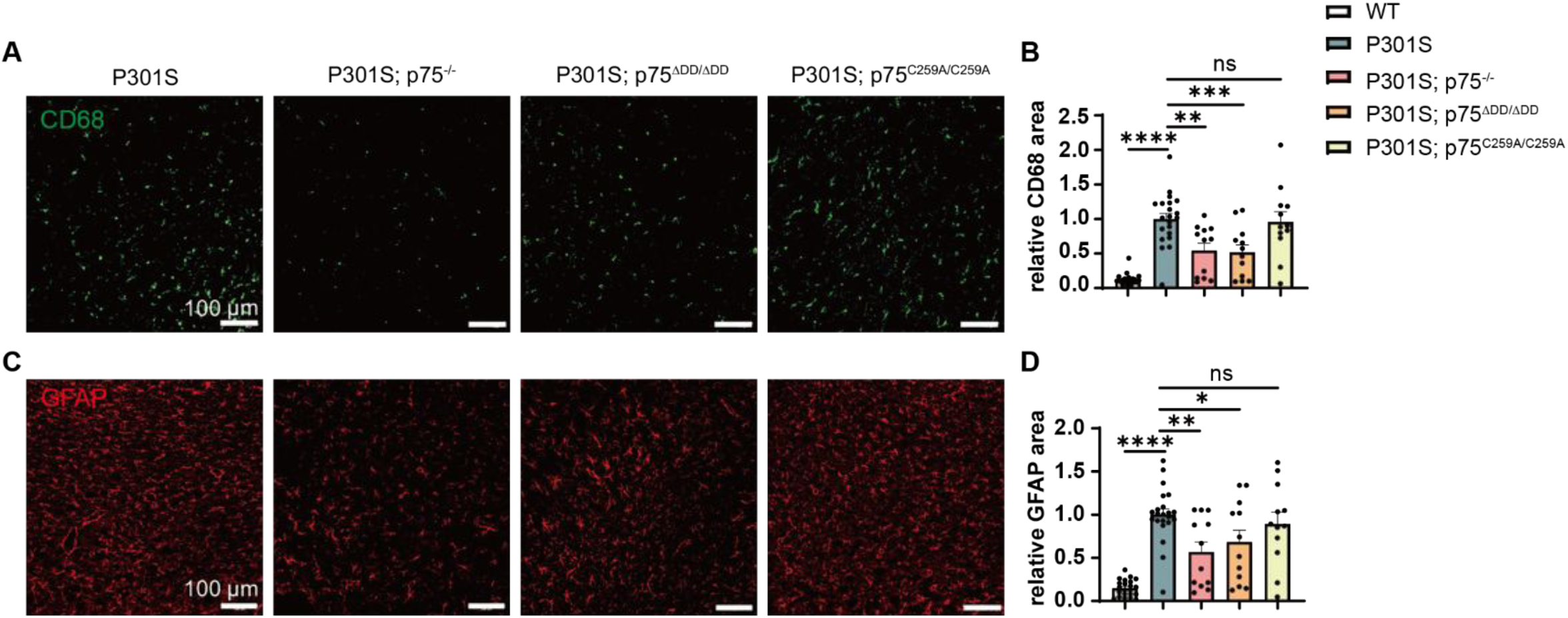
Reduced microgliosis and astrogliosis in the PIR of P301S mice lacking death domain and global knock-out but not transmembrane Cys_259_ of p75^NTR^. A) Representative microgliosis confocal images of CD68 (green) immunohistology in the PIR of 9-month-old WT, P301S, P301S; p75^-/-^, P301S; p75^ΔDD/ΔDD^ and P301S; p75^C259A/C259A^, as indicated. Scale bar, 100 μm. B) Quantification of the level of microgliosis assessed by the relative CD68 area. N=22(WT, P301S), 12(P301S; p75^-/-^, P301S; p75^ΔDD/ΔDD^, P301S; p75^C259A/C259A^) mice per genotype, respectively. One-way ANOVA followed by Dunnett’s multiple comparisons test, mean ± SEM. **p<0.01, ***p<0.001, ****p<0.0001. C) Representative microgliosis confocal images of GFAP (red) immunohistology in the PIR of 9-month-old WT, P301S, P301S; p75^-/-^, P301S; p75^ΔDD/ΔDD^ and P301S; p75^C259A/C259A^, as indicated. Scale bar, 100 μm. D) Quantification of the level of microgliosis assessed by the relative GFAP area. N=22(WT, P301S), 12(P301S; p75^-/-^, P301S; p75^ΔDD/ΔDD^, P301S; p75^C259A/C259A^) mice per genotype, respectively. One-way ANOVA followed by Dunnett’s multiple comparisons test, mean ± SEM. *p<0.05, **p<0.01, ****p<0.0001.

**Figure S4.**
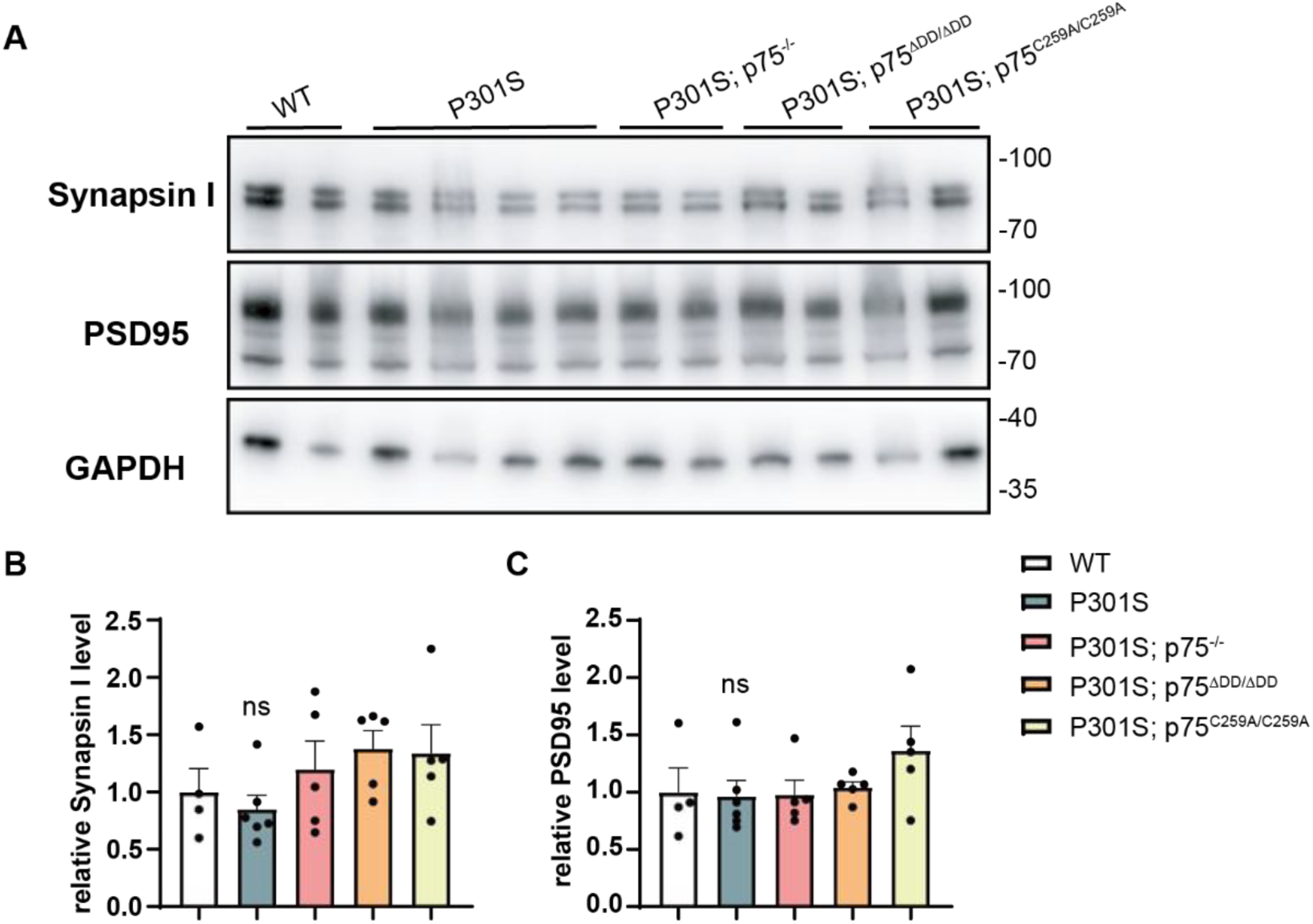
Comparable synaptic protein expression level among WT and transgenic mouse models. A) Representative Western blot gels of whole hippocampus extract from 9-month-old WT, P301S, P301S; p75^-/-^, P301S; p75^ΔDD/ΔDD^ and P301S; p75^C259A/C259A^ as indicated using Synapsin I and PSD95 antibody. Molecular weights are indicated in kDa. B-C) Quantification of Synapsin I (B) and PSD95 (C) protein level from the whole tissue extract relative to GAPDH. N=4(WT), 6(P301S), 5(P301S; p75^-/-^, P301S; p75^ΔDD/ΔDD^, P301S; p75^C259A/C259A^) mice per genotype, respectively. One-way ANOVA followed by Dunnett’s multiple comparisons test, mean ± SEM. No significant difference between WT, P301S; p75^-/-^, P301S; p75^ΔDD/ΔDD^, P301S; p75^C259A/C259A^ versus P301S, respectively.

**Fig S5.**
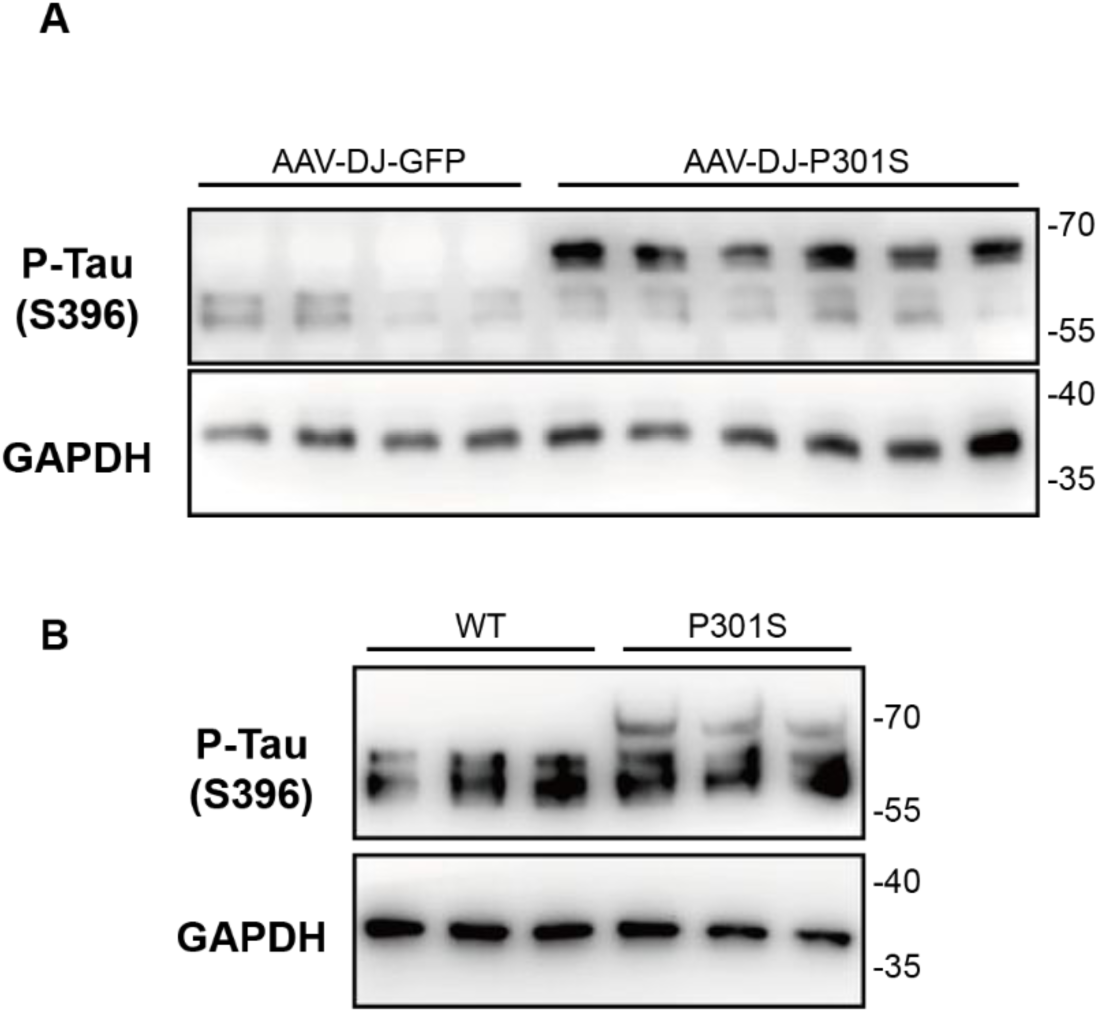
Primary cerebral cortical neurons infected with AAV-DJ-P301S virus or expressing P301S transgene have hyperphosphorylated Tau expression. A) Representative Western blot gels of the WT cerebral cortical neuron infected with either AAV-DJ-GFP or -P301S virus, as indicated. Molecular weights are indicated in kDa. B) Representative Western blot gels of the cerebral cortical neuron from WT and P301S embryos, as indicated. Molecular weights are indicated in kDa.

**Fig S6.**
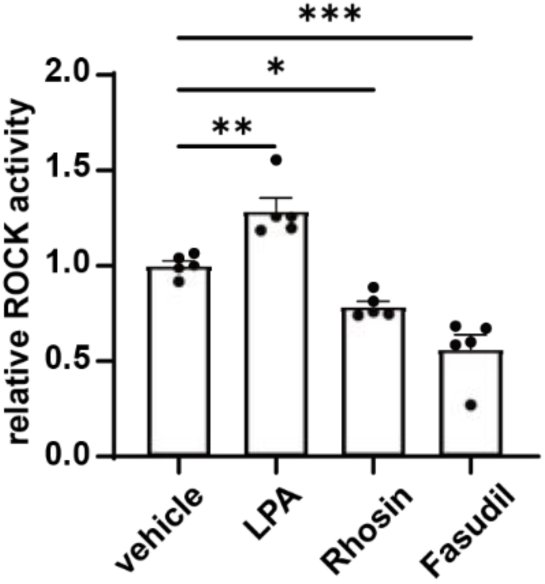
Drugs targeting at RhoA could regulate ROCK activity in primary cultured cerebral cortical neuron. Quantification results of ROCK ELISA from primary cultured cerebral cortical neuron from P301S embryos. Results shown are from 5 independent experiments with 2 replicates each. One-way ANOVA followed by Dunnett’s multiple comparisons test, mean ± SEM. *p<0.05, **p<0.01, ***p<0.001.

**Fig S7.**
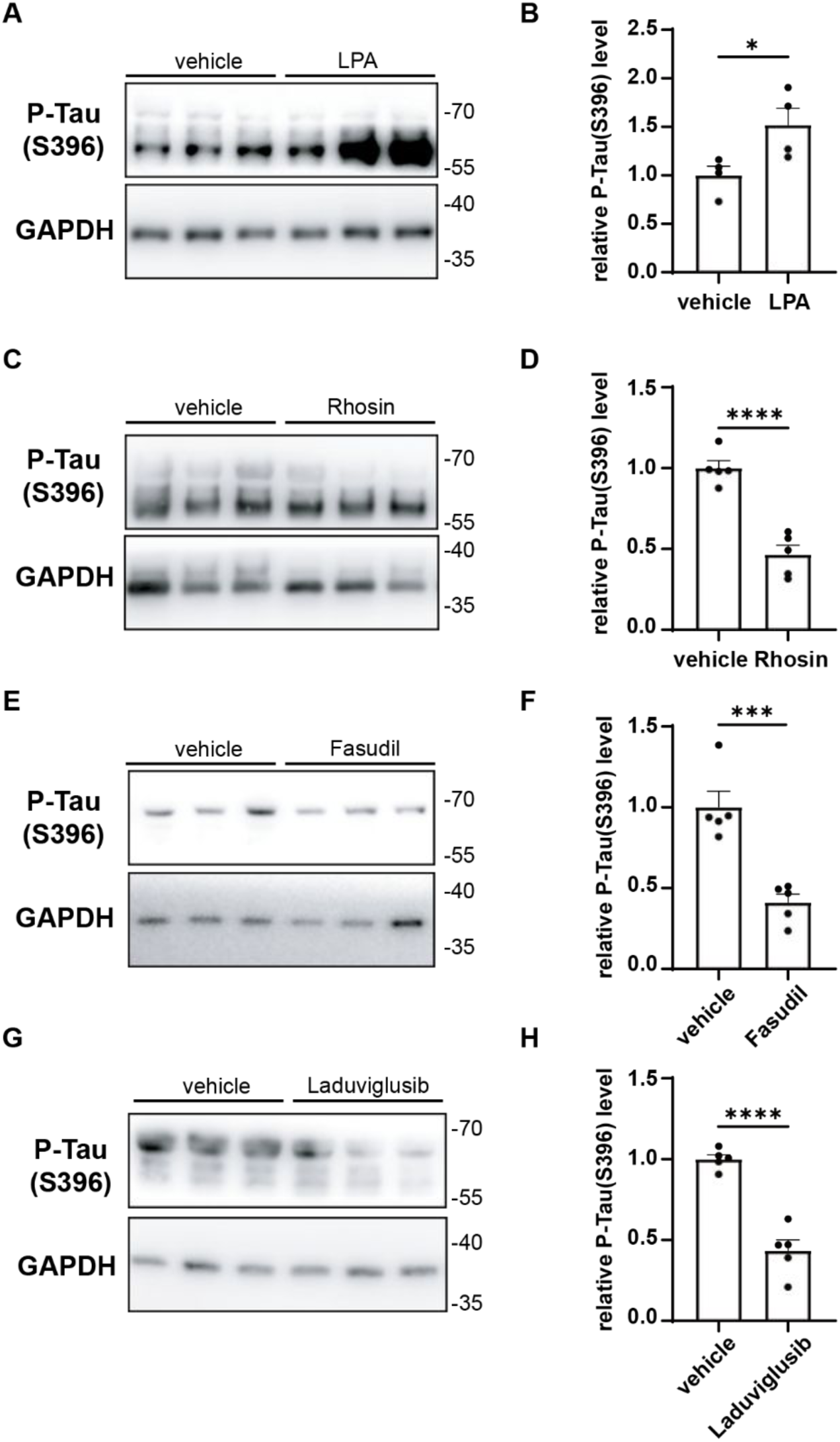
Drugs targeting at RhoA, ROCK and GSK3β could regulate P-Tau level in primary cultured cerebral cortical neuron. A) Representative Western blot gels of the WT cerebral cortical neuron infected with AAV-DJ-P301S virus and treated with either vehicle or LPA (1 μM) for 24 h, as indicated. Molecular weights are indicated in kDa. B) Quantification results of Western blot from WT cerebral cortical neurons infected with AAV-DJ-P301S virus. Results shown are from 4 independent experiments with 3 replicates each. Unpaired Student’s t-test, mean ± SEM. *p<0.05. C) Representative Western blot gels of the WT cerebral cortical neuron infected with AAV-DJ-P301S virus and treated with either vehicle or Rhosin (10 μM) for 24 h, as indicated. Molecular weights are indicated in kDa. D) Quantification results of Western blot from WT cerebral cortical neurons infected with AAV-DJ-P301S virus. Results shown are from 5 independent experiments with 3 replicates each. Unpaired Student’s t-test, mean ± SEM. ****p<0.0001. E) Representative Western blot gels of the WT cerebral cortical neuron infected with AAV-DJ-P301S virus and treated with either vehicle or Fasudil (15 μg/mL) for 24 h, as indicated. Molecular weights are indicated in kDa. F) Quantification results of Western blot from WT cerebral cortical neurons infected with AAV-DJ-P301S virus. Results shown are from 5 independent experiments with 3 replicates each. Unpaired Student’s t-test, mean ± SEM. ***p<0.001. G) Representative Western blot gels of the WT cerebral cortical neuron infected with AAV-DJ-P301S virus and treated with either vehicle or Laduviglusib (10 μM) for 24 h, as indicated. Molecular weights are indicated in kDa. H) Quantification results of Western blot from WT cerebral cortical neurons infected with AAV-DJ-P301S virus. Results shown are from 5 independent experiments with 3 replicates each. Unpaired Student’s t-test, mean ± SEM. ****p<0.0001.

**Fig S8.**
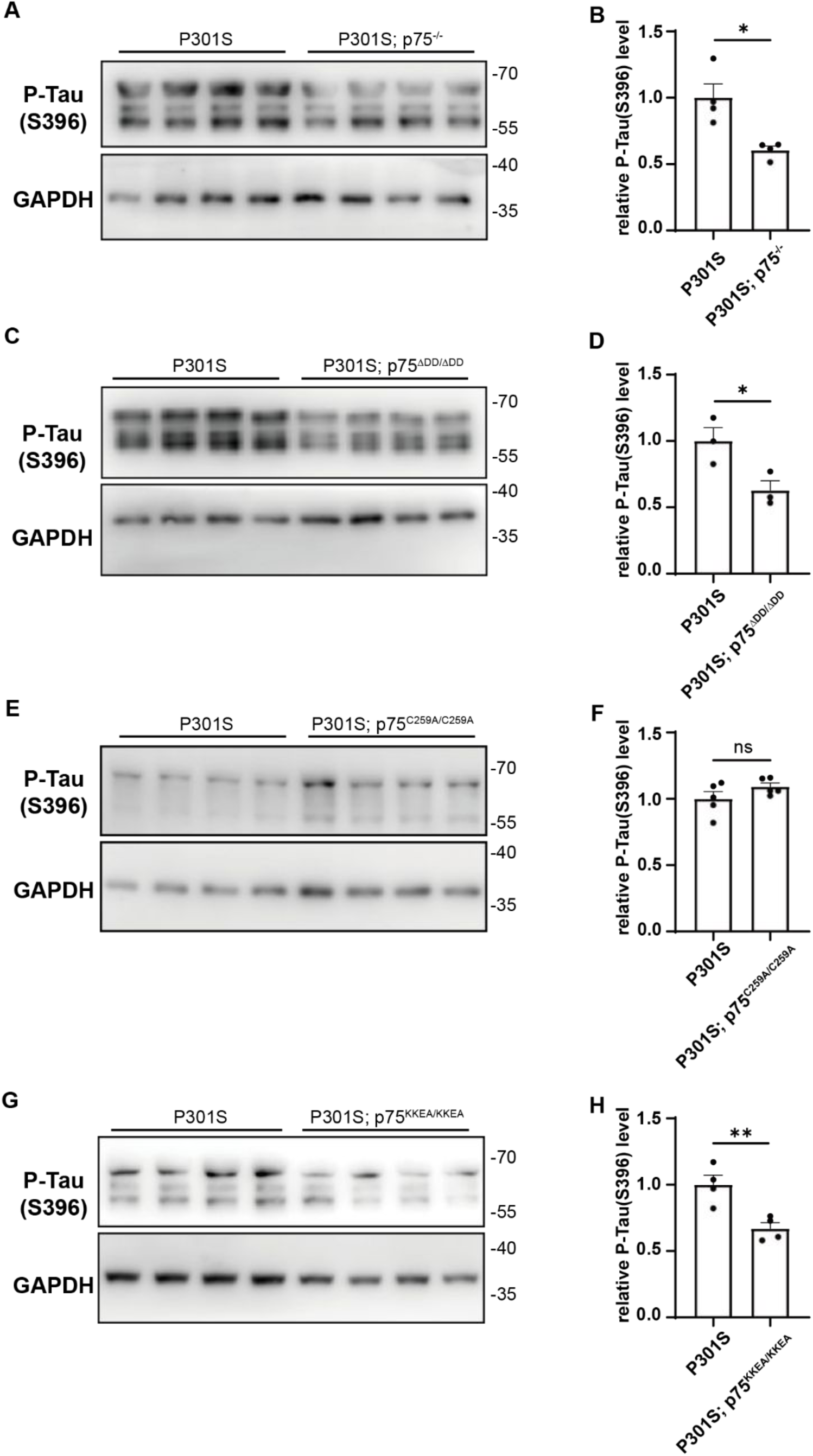
Reduced P-Tau level in the primary cultured hippocampal neuron of P301S mice lacking death domain, global knock-out and RhoA-deficient but not transmembrane Cys_259_ of p75^NTR^. A) Representative Western blot gels of the neuron from P301S and P301S; p75^-/-^ embryos, as indicated. Molecular weights are indicated in kDa. B) Quantification results of Western blot from P301S and P301S; p75^-/-^ neurons. Results shown are from 4 independent experiments with 4 replicates from each. Unpaired Student’s t-test, mean ± SEM. *p<0.05. C) Representative Western blot gels of the neuron from P301S and P301S; p75^ΔDD/ΔDD^ embryos, as indicated. D) Quantification results of Western blot from P301S and P301S; p75^ΔDD/ΔDD^ neurons. Results shown are from 3 independent experiments with 4 replicates from each. Unpaired Student’s t-test, mean ± SEM. *p<0.05. E) Representative Western blot gels of the neuron from P301S and P301S; p75^C259A/C259A^ embryos, as indicated. F) Quantification results of Western blot from P301S and P301S; p75^C259A/C259A^ neurons. Results shown are from 5 independent experiments with 4 replicates from each. Unpaired Student’s t-test, mean ± SEM, ns, no significance. G) Representative Western blot gels of the neuron from P301S and P301S; p75^KKEA/KKEA^ embryos, as indicated. H) Quantification results of Western blot from P301S and P301S; p75^KKEA/KKEA^ neurons. Results shown are from 4 independent experiments with 4 replicates from each. Unpaired Student’s t-test, mean ± SEM. **p<0.01.

**Fig S9.**
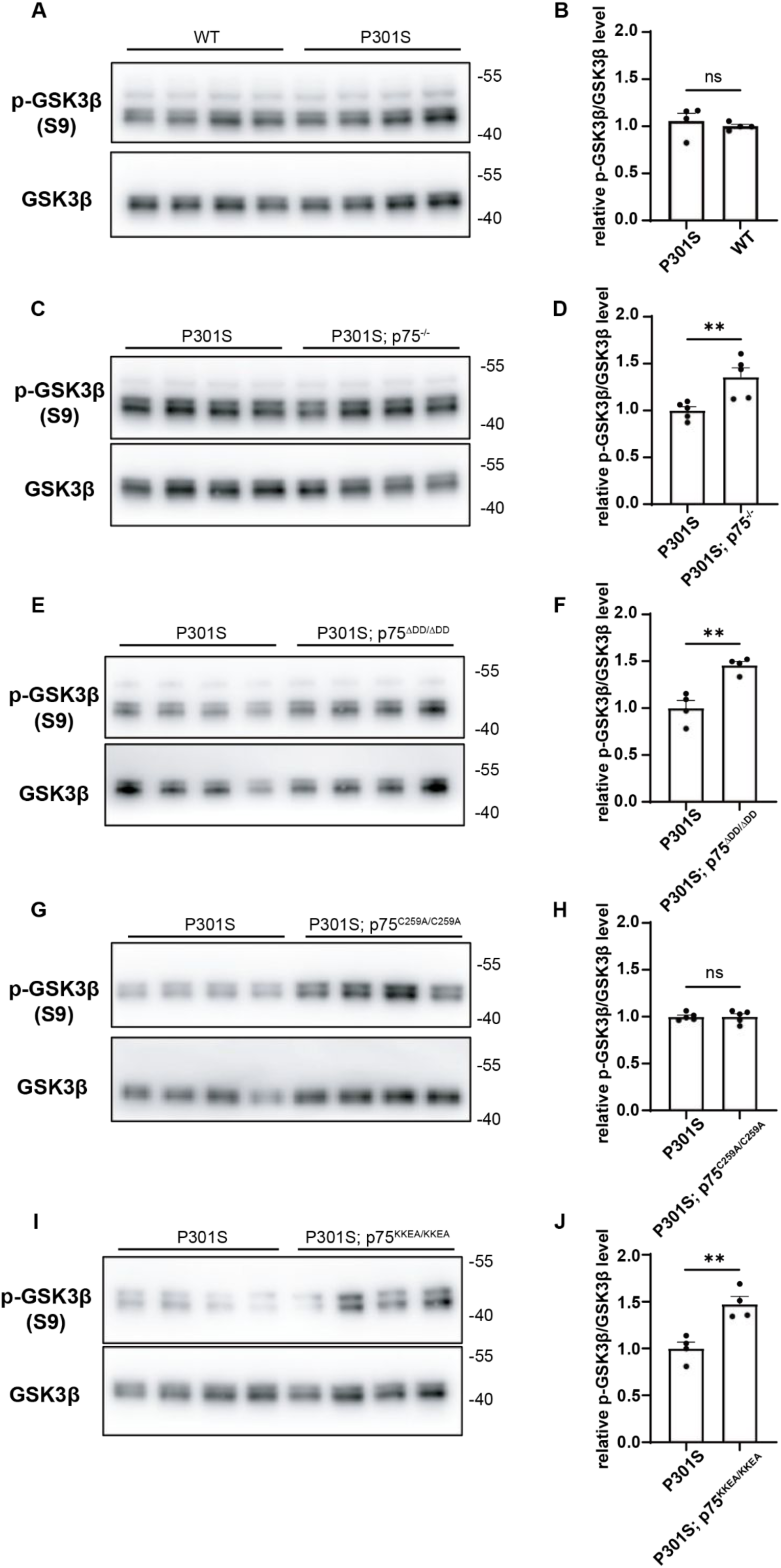
Reduced GSK3β activity in the primary cultured cerebral cortical neuron of P301S mice lacking death domain, global knock-out and RhoA-deficient but not transmembrane Cys_259_ of p75^NTR^. A) Representative Western blot gels of the neuron from P301S and WT embryos as indicated using S9 antibody relative to GSK3β. Molecular weights are indicated in kDa. B) Quantification results of Western blot from P301S and WT neurons. Results shown are from 4 independent experiments with 4 replicates each. Unpaired Student’s t-test, mean ± SEM. No significant difference between WT versus P301S. C) Representative Western blot gels of the neuron from P301S and P301S; p75^-/-^ embryos as indicated using S9 antibody relative to GSK3β. Molecular weights are indicated in kDa. D) Quantification results of Western blot from P301S and P301S; p75^-/-^ neurons. Results shown are from 4 independent experiments with 5 replicates each. Unpaired Student’s t-test, mean ± SEM. **p<0.01. E) Representative Western blot gels of the neuron from P301S and P301S; p75^ΔDD/ΔDD^ embryos, as indicated. F) Quantification results of Western blot from P301S and P301S; p75^ΔDD/ΔDD^ neurons. Results shown are from 4 independent experiments with 4 replicates from each. Unpaired Student’s t-test, mean ± SEM. **p<0.01. G) Representative Western blot gels of the neuron from P301S and P301S; p75^C259A/C259A^ embryos, as indicated. H) Quantification results of Western blot from P301S and P301S; p75^C259A/C259A^ neurons. Results shown are from 4 independent experiments with 5 replicates each. Unpaired Student’s t-test, mean ± SEM, ns, no significance. I) Representative Western blot gels of the neuron from P301S and P301S; p75^KKEA/KKEA^ embryos, as indicated. J) Quantification results of Western blot from P301S and P301S; p75^KKEA/KKEA^ neurons. Results shown are from 4 independent experiments with 4 replicates each. Unpaired Student’s t-test, mean ± SEM. **p<0.01.

## Notes

### Competing Interest Statement

The authors have declared no competing interest.

